# Distinct roles of Hsp110 proteins in regulating prion propagation

**DOI:** 10.64898/2026.10.14.758948

**Authors:** Matteo Mosca, Mariam Ansari, Cristóbal Marrero-Winkens, Charu Sankaran, Basant Abdulrahman, Hermann Schatzl

**Author notes:** This work is an equal contribution (Matteo Mosca, Mariam Ansari).

## Abstract

Prion diseases are fatal, transmissible neurodegenerative disorders characterized by the conformational conversion of the cellular prion protein (PrP^C)^ into its pathogenic isoform PrP^Sc^. Propagation of prions depends on the fragmentation of PrP^Sc^ aggregates to generate new seeds. However, the factors regulating this process remain incompletely understood. For yeast prions, Hsp104 that has no mammalian homologue, is firmly established as a disaggregase. Hsp110 has emerged as a potential mediator of protein disaggregation and aggregate remodeling in mammalian cells. Although Hsp110 proteins have been implicated in several protein misfolding disorders, their individual contributions to mammalian prion propagation have not been characterized. In this study, we examined the roles of the three Hsp110 proteins—Apg-1, Apg-2, and Hsp105—in cellular prion infection. CRISPR-generated knockout cell lines were established for each Hsp110 and evaluated for their effect on *de novo* prion infection. PrP^Sc^ accumulation was assessed by immunoblot across serial passages and up to 40 days, and prion seeding activity was quantified using real-time quaking-induced conversion (RT-QuIC). Loss of individual Hsp110’s produced distinct effects on prion propagation. Apg-1-deficient cells exhibited strongly reduced levels of PrP^Sc^ over all passages analyzed, suggesting a role for Apg-1 in establishing persistent prion infection. Interestingly, prion seeding activity was detectable in later passages. Hsp105-deficient cells displayed reduced PrP^Sc^ levels during the early stages, indicating that Hsp105 contributes to early prion replication. In contrast, Apg-2 knockout resulted in high prion seeding activity over all passages analyzed and only slightly reduced PrP^Sc^ levels in early passages. Collectively, these findings demonstrate that Hsp110 family members perform distinct functions during early phases of prion infection and establishment of persistent infection. This work provides new insight into the role of molecular chaperones in prion biology and identifies Hsp110 proteins as potential modulators of prion replication and potentially therapeutic targets for protein misfolding diseases.

## Introduction

Prion diseases are unique neurodegenerative disorders caused by the misfolding of the endogenous cellular prion protein (PrP^C^) [1]. They are not only infectious but also fatal and, unlike most transmissible diseases, do not involve nucleic acids but rather utilize proteins for spreading [1]. The misfolded form of the prion protein (PrP^Sc^) is prone to aggregation and causes lesions to the central nervous system, including neuronal loss, vacuolation and astrogliosis [2]. Various prion diseases exist in both animals and humans, and the potential for zoonotic transmission coupled with the lack of treatment or prophylaxis results in this disease being of concern. The exact mechanism of prion protein conversion remains unknown; however, it is established that interactions take place between the cellular and misfolded forms, resulting in the misfolding of the cellular form. In the seeded nucleation model, it is hypothesized that individual PrP^Sc^ molecules nucleate to form small seeds that work to recruit and convert PrP^C^ to PrP^Sc^ on the ends. The seeds oligomerize and elongate to form protofibrils, which then begin to aggregate to build dense fibrils [3]. After reaching a certain size, further conversion of PrP^C^ becomes inefficient, so there is a postulated step of fragmentation that breaks down protofibrils and fibrils back into seeds [3, 4].

Protein misfolding is not unique to disease states and occurs at a small scale under physiological conditions [5]. Cells can respond to protein misfolding through protein refolding or protein degradation. Since prion diseases involve protein misfolding, cellular protein refolding and quality control machineries have become a pathway of interest. Our group and others have proposed that these pathways may be involved in mammalian prion fragmentation and work to break down prion fibrils [3, 6]. One such refolding machinery is the Hsp70 chaperone system which is comprised of three main components: Hsp70, a J-domain protein (JDP) and Hsp110 [7]. Hsp70 utilizes a binding and release mechanism to promiscuously bind exposed hydrophobic regions and targets both misfolded proteins and nascent unfolded proteins [7, 8]. Hsp70 contains two domains: a substrate-binding domain (SBD) that binds the misfolded protein and a nucleotide-binding domain (NBD), and requires two chaperones, a JDP and Hsp110, for optimal function [7]. The JDP captures nascent proteins and recruits other substrates to Hsp70 to form an Hsp70-JDP complex, whereas Hsp110 acts as a nucleotide exchange factor [9]. There are three homologues of Hsp110, including Apg-1, Apg-2 and Hsp105 [7, 10]. Besides the nucleotide exchange factor activity, Hsp110 has been implicated in protein refolding and disaggregation [11]. Hsp110 was shown to work alongside Hsp70 and JDP to dissolve large aggregates and recover native protein structure in cell-free systems [12]. Immunodepletion of Hsp105 and Apg-2 showed slightly reduced and greatly reduced disaggregase activity, respectively [12].

In yeast, the Hsp70-Hsp40-Hsp104 tri-chaperone system has been shown to disaggregate amorphous protein aggregates, allowing for refolding [13]. Furthermore, Hsp104 has been shown to be required for the propagation of yeast prions [14]. There is no mammalian homologue for Hsp104, and as such, studies are underway to identify a functional equivalent, with the Hsp70 system, including Hsp110, being a potential candidate [12]. Moreover, although the Hsp110 family members work to break down aggregates, their role in neurodegenerative diseases is still under debate, with two main standpoints. First, it is hypothesized that the Hsp70 system breaks down large aggregates to produce seeds to promote prion propagation. Tittelmeier and colleagues showed this through *in vivo* experiments with *C. elegans*, where Hsp110 was transiently knocked down, causing a reduction in the number of α-synuclein foci and overall reduced toxicity, supporting the notion that Hsp110 promotes prion spreading through seed propagation [15]. Similarly, it has been shown that depletion of Hsp110 decreased amyloid beta aggregation in a *C. elegans* model [16]. The opposing view states that the Hsp70 system works to clear prion aggregates, resulting in a reduction in the detrimental effects of the accumulation of prion fibrils. This was shown in studies that analyzed Hsp70 expression *in vivo* in Drosophila expressing mammalian PrP [6]. This study showed a reduction in the expression of the components of the Hsp70 system – Hsp70, Hsp110 and JDP – in prion-exposed Drosophila [6]. Moreover, their results showed that downregulation of Hsp110 or JDP increased toxicity, whereas overexpression of these chaperones reduced toxicity with increasing doses of prion inoculum [6]. This is supported by work from Eroglu et al. that showed that depletion of Hsp110 also increased tau aggregation in a mouse model [17]. Overall, Hsp110 modulation has been shown to affect protein aggregation in a variety of neurodegenerative models.

With respect to the effects of each Hsp110 homologue, Hsp105 was first observed to prevent firefly luciferase aggregation [18]. Hsp105 has been observed to play a role in protein disaggregation in various experimental models of protein misfolding diseases including suppression of SOD1 aggregation associated with ALS [19], suppression of poly-Q aggregation [20], stabilization of CFTR (cystic fibrosis transmembrane conductance regulator) aggregation in epithelial cells associated with cystic fibrosis [21], and inhibition of amyloid Aβ-42 and α-synuclein fibrils *in vitro* in Drosophila [22]. Furthermore, more pronounced disaggregrase effects were observed when Apg-2 was depleted in comparison to Hsp105, suggesting a greater role of Apg-2 in disaggregation [12]. Apg-2 was later shown to slowly disassemble α-synuclein aggregates using a depolymerization model where chaperones work to remove monomers from the ends of fibrils to slowly solubilize them, thus supporting chaperone-mediated fragmentation [23–26]. It has been reported that monomers produced by Hsp70 machinery breaking down α-synuclein fibrils were soluble and less toxic than α-synuclein amyloid fibrils [25]. On the other hand, most recently, Shoup & Priola showed that Hsp105 and Apg-2 reduced the levels of purified PrP^Sc^ in cell-free *in vitro* assays for two mouse-adapted prion strains, 22L and 87V [27]. Their work showed the formation of seeding activity when 22L prion fibrils were disaggregated *in vitro* using Apg-2 or Hsp105 [27]. Lastly, current research on the role of Apg-1 in prion disease is absent; however, it has been implicated in a neuroprotective role in various protein misfolding diseases, suppressing the formation of amyloid amyloid beta, α-synuclein, and poly-glutamine aggregates [22, 28], as well as increasing the lifespan of mice with an ALS-associated SOD1 mutation overexpressing Apg-1 [29]. All this suggests that Hsp110s act as modifiers of aggregation, cellular toxicity and wider neurodegeneration as seen in protein-misfolding disorders.

There is currently no research that compares the effects of each of the three Hsp110 homologues with respect to prion propagation. In this study, using a murine neuroblastoma N2a cell culture model, we generated single knockout cell lines to assess the roles of each Hsp110 homologue in prion infection. These single knockout cell lines were assessed for cell viability and then utilized in *de novo* infection experiments using 22L mouse-adapted prions and cells were monitored for up to 40 days after infection. Our results showed that knockout of Apg-1, Apg-2 and Hsp105 impacts both PrP^Sc^ accumulation and prion seeding activity in *de novo* infection experiments. Apg-1KO cells showed strongly reduced levels of PrP^Sc^ over all passages analyzed, whereas Hsp105KO cells showed lower levels of PrP^Sc^ in early passages of infection. Apg-2KO cells resulted in high prion seeding activity over all passages analyzed and only slightly reduced PrP^Sc^ levels in early passages. RT-QuIC experiments also highlighted distinct, cell-line specific effects across passages. Overall, our data demonstrate that individual Hsp110 members have distinct regulatory roles in prion infection and propagation.

## Materials and Methods

### Ethics statement

All C57/Bl6 mice used in the experiments involving infection with 22L, RML and Me7 prions were handled and euthanized according to the approved guidelines and regulations of the University of Calgary Health Science Animal Care Committee (HSACC) under protocol number AC18-0047 and in strict accordance with the Canadian Council for Animal Care (CCAC) guidelines. The mice were housed in an environment with controlled temperature and humidity with alternating light and dark cycles of 12 hours. Mice were given free access to food and drinking water until the time of euthanasia.

### Cell culture

Murine neuroblastoma cells (N2a) subclone K-21 derived from the mouse neuroblastoma cell line previously described in Gilch et al., 2001 [30] and Abdulrahman et al., 2018 [31] were used for the duration of this project. Murine catecholaminergic neuronal (CAD5) cells were established and gifted by Dr. S. Mahal and Dr. C. Weissmann (Scripps Research, FL, USA). Murine embryonic fibroblasts (MEF) cells were kindly provided by Dr. N. Mizushima (Tokyo Medical and Dental University, Japan). N2a and CAD5 cells were cultured using OptiMEM + GlutaMAX media (Gibco, #51985034), supplemented with 10% bovine growth serum (Hyclone, #SH30541.03), filtered with a 0.45 μm filter, and 1% penicillin/streptomycin (10000 U/mL pen/strep; Gibco, #15140122). Cells were grown to 80-100% confluency in 10-cm cell culture plates and passaged 1:10 or 1:20 using 1 mL trypsin-EDTA after a 10 mL wash with 1x PBS, pH 7.4, (Gibco, #70011044). Cells were incubated at 37°C with 5% CO_2_ and reached confluency in 4-5 days. MEF cells were cultured under the same conditions but with DMEM media (Gibco, #11965092), supplemented with 10% fetal bovine serum (Sigma-Aldrich, #F2442) and 1% pen/strep.

### Transient knockdown

Proprietary siRNAs were obtained from Dharmacon (Lafayette, CO, USA) as pools of four distinct siRNA sequences directed against the same target. These siRNAs included: Murine Apg-1 *(Hspa4l)* (#L-041742-01), Murine Apg-2 (Hspa4) (#L-062183-01), Murine Hsp105 *(Hsph1)* (#L-004972-00) and non-targeting control siRNA (#D-001810-10).

For the knockdown, 350,000 cells were seeded into 6-cm cell culture plates and grown for one day. The next day, 200 μl of 25 nM siRNA solution was prepared in media and incubated at room temperature for 5 minutes and then added dropwise to 200 μl of 12.35 ng/μl solution of Lipofectamine 2000 (Invitrogen, #11668027) prepared in media. The siRNA-Lipofectamine 2000 mixture was incubated for 30 minutes at room temperature, then added dropwise to the cells along with 1.6 ml of complete antibiotic-free media. Cells were incubated overnight at 37°C, and the next day, media was replaced with 2 ml of fresh complete media. Knockdown was confirmed with immunoblotting.

### Transient overexpression

For transient overexpression, 300,000 cells were seeded into 6-cm cell culture plates and grown for one day. The next day, 200 μl of media, 1.5 μl PLUS Reagent (Invitrogen, #A12621) and 1 μg of plasmid DNA was prepared and incubated at room temperature for 5 minutes and then added dropwise to 200 μl of media and 3 μl of Lipofectamine LTX (Invitrogen, #A12621). The DNA-Plus Reagent-Lipofectamine LTX mixture was incubated at room temperature for 20 minutes, then added dropwise to the cell culture plates with complete media. Cells were incubated overnight at 37°C, and the next day, the media was replaced with 2 ml of fresh complete media. Overexpression was confirmed with immunoblotting.

### Knockout cell lines

#### Hsp105 (Hsph1) knockout in N2a cells

The following constructs targeting exons 1 and 7 of the Hsp105 (*Hsph1*) gene were used: Exon 1: target ID: Hsp105-1, target sequence: 5’-TACATTGCGGTGGCGCGGGC −3’; Exon 7: target ID: Hsp105-7-a, target sequence: 5’-GCTTTTGATCCCTTCTTAGG −3’; Exon 7: target ID: Hsp105-7-b, target sequence: 5’-AAATTCGAGCCCTCCTTCGTC −3’.

The pSpCas9(BB)-2A-GFP (PX458) plasmid vector encoding *Streptococcus pyogenes* Cas9 enzyme with a cloning backbone for single guide RNAs (sgRNAs) was a kind gift from Dr. Feng Zhang (Watertown, MA, USA, Addgene plasmid # 48138) and was used for the generation of the *Hsph1* knock-out cell line. All guide RNAs (gRNAs) were designed using Benchling software (San Francisco, CA, USA) and procured from the University of Calgary Core DNA facility. gRNA sequences were cloned into the PX458 backbone according to published protocols [32]. Briefly, the plasmid was digested and dephosphorylated using FastDigest Bpil from ThermoFisher (Waltham, MA, USA) and Quick CIP from NEB (Ipswich, MA, USA) for 30 min at 37°C, followed by visualization and purification on an agarose gel. Complementary oligonucleotide pairs encoding the gRNA sequence were phosphorylated and annealed using T4 PNK from NEB (Ipswich, MA, USA), in a thermocycler at 37°C for 30 min, 95°C for 5 min and then ramped down to 25°C at 5°/min. Next, the ligation reaction was established using 50 ng of BbsI-digested PX485 plasmid, 1 μl of the phosphorylated and annealed oligo duplex reaction and Quick Ligase from NEB (Ipswich, MA, USA). The reaction was incubated at RT for 1 h, followed by transformation into Stbl3 *Escherichia coli* from ThermoFisher (Waltham, MA, USA).

#### Apg-1 (Hspa4l) and Apg-2 (Hspa4) knockout in N2a cells

Guide RNAs were designed and supplied by Synthego Gene Knockout kits (GKO Kits) for Apg-1 and Apg-2 that included a mixture of 3 gRNAs. The following constructs targeting exon 1 of the Apg-1 (*Hspa4l*) gene and exon 1 of the Apg 2 (*Hspa4*) gene were used: Exon 1: (Apg 1) gRNA sequence 1: 5’-GGCGUCCCGGGCCCCACCCC −3’; Exon 1: (Apg 1) gRNA sequence 2: 5’-GCACCUGUCGCUGUACUCGU-3’; Exon 1: (Apg 1) gRNA sequence 3: 5’-CUACAGCGAUGUAGCAGUUG −3’; Exon 1: (Apg 2) gRNA sequence 1: 5’-GUCGCUGUGGCCCGCGCCGG-3’; Exon 1: (Apg 2) gRNA sequence 2: 5’-GCCCCACUUACGGCGUGCAG-3’; Exon 1: (Apg 2) gRNA sequence 3: 5’-GCAGGCCCAGGAUCCGAAGC-3’;

The protocol from the Synthego manufacturer was used for the CRISPR knockout experiments for Apg-1 and Apg-2. 100,000 wild-type N2a cells were grown and spun at 200 g for 5 minutes. The supernatant was removed and 500 μl of serum-free media was added. 25 μl of serum-free media was added to 1.3 μl of 3 μM gRNA, 1 μl of 3 μM Cas9 and 1 μl of Lipofectamine Cas9 PLUS reagent (Invitrogen, #CMAX00003) and incubated for 5 minutes. In a separate tube, 25 μl of serum-free media was added to 1.5 μl of CRISPRMAX transfection reagent (Invitrogen, #CMAX00003). After the incubation, the CRISPRMAX mix was added to the gRNA-Cas9 mix and vortexed. After a 10 min incubation, the RNP-transfection reagents were added to the cell suspension and plated into a 24-well cell culture plate with extra media. Cells were monitored, and the following day, the wells were replaced with fresh serum-free media. Mixed populations of cells were plated for limiting dilution single-cell cloning by diluting cell suspensions to 8 cells/ml. 200 μl of cell suspension was added to each well of 96-well plates and monitored for 4-6 weeks. Wells with single-cell colonies were passaged to 24-well plates, then 6-well plates and finally 10-cm plates before being tested with immunoblot and DNA sequencing to confirm knockout.

### DNA extraction, PCR and sequencing

Genomic DNA was extracted from single-cell clones and used as a template for amplification of fragments for knockout confirmation. The following primers were used: Apg-1 Exon 1, forward primer: 5’-GGGTCCTCTCCTTTGCTTCT-3’, reverse primer: AAGCAGAGATGAGGCGAGC-3’; Apg-2 Exon 1, forward primer: 5’-GGTAGAGGCCCCCGTCTTTC-3’, reverse primer: 5’-GTCCATCCCACGGTCTCAGG-3’; Hsp105 Exon 7, forward primer: 5’-GCAGGTCCGTGCTGTTAGAA-3’, reverse primer: 5’-ACCAACCTTAGGTTCTAGGCAC-3’. DNA was extracted from cells using the E.Z.N.A. Tissue Culture Kit (OMEGA Bio-Tek, #D3396-00S) following its protocol. Confluent cells were trypsinized and washed with PBS, then lysed using the protease solution provided in the kit. RNase A was also added during this step. The solution was then heated, centrifuged and filtered after adding various buffers from the kit. DNA was eluted using 100μl ultrapure distilled water and stored at 4°C for short-term storage and −20°C for long-term storage. DNA samples were diluted to 50-100 μg/μl for PCR and other analyses. Genomic DNA was amplified using PCR before bacterial cloning. PCR reaction conditions were as follows: 1 μl of 50-100 µg/μl template DNA, 1 μl of 10 μM forward primer, 1 μl of 10 μM reverse primer, 5 μl of 10x Pfu buffer (Agilent, #600153), 1 μl of 10 μM dNTP (Invitrogen, #10297018), 0.5 μl of 2.5 U/μl Pfu polymerase (Agilent, #600153) and nuclease-free water in a total reaction volume of 50 μl. For Apg-1 and Apg-2 genomic DNA amplification, the PCR reaction mix had the addition of 1.5 μl of DMSO (Sigma-Aldrich, #472301). Conditions for the PCR were 2 minutes of initial denaturation at 95°C, followed by 35 cycles of denaturation at 95°C for 20 seconds, annealing at differing T_m_ values (57°C for Hsp105 and 55°C for Apg-1 and Apg-2) for 30 seconds, and extension at 72°C for 15 seconds, with a final extension at 72°C for 3 minutes. All PCR products were purified and visualized by agarose gel electrophoresis and then amplified using the Zero Blunt TOPO PCR Cloning Kit (ThermoFisher, #450245). DNA fragments were cloned into One Shot TOP10 Chemically Competent *E. coli* (ThermoFisher, #C404010) and kanamycin was used as a selection marker. Purified plasmid DNA was sent to the University of Calgary Sanger DNA Sequencing facility with M13 primers to confirm the knockouts.

### LDH assay

Lactate dehydrogenase assays were performed using the Cytotoxicity Detection kit (Roche, #11644793001) following its protocol. 70 μl of cell culture supernatant was added to 96-well plates. 250 μl of Reagent 1 was mixed with 11.25 ml of Reagent 2 and added to each well of supernatant with a volume of 70 μl. The cell culture plate was kept in darkness for 5 minutes before stopping the reaction with the addition of 30 μl of 1 N HCl in each well. The absorbance values of the plate were measured at 490 nm and 650 nm as a reference. Supernatant from cells treated with 10% TritonX-100 (ThermoFisher, #A16046.AE) was used as a positive control.

### XTT assays

XTT assays were performed using the Cell Proliferation Kit II (Roche, # 11465015001). 7,000 cells were seeded in 100 μl culture media per well in 96-well cell culture plates and incubated overnight. The following day 100 μl of XTT activator was added to 5 ml of XTT reagent and 50 μl of this mixture was added to each well. The plate was incubated for 3 hours at 37°C with 5% CO_2._ The absorbance values of the plate were measured at 490 nm and 650 nm as a reference. Cells treated with Triton-X100 were used as a negative control.

### Cell lysis for uninfected cells

Uninfected cells were lysed with 1X lysis buffer (10 mM Tris-HCl, pH 7.5; 100 mM NaCl; 10 mM EDTA; 0.5% Triton X-100; 0.5% sodium deoxycholate) with Roche’s cOmplete protease inhibitor cocktail (1 tablet in 10 ml of cell lysis buffer). Lysate was added to a 1.5 ml microcentrifuge tube and centrifuged at 15,000 g for 1 minute and the supernatant was precipitated with excess methanol (1 ml of lysate with 5 ml methanol), and stored overnight at −20°C, or for one hour at −80°C. After incubation, lysates were centrifuged at 3,500 rpm for 25 minutes, and the protein pellet was resuspended with 200 μl of TNE buffer (50 mM Tris-HCl pH 7.5, 150 mM NaCl, 5 mM EDTA) and 100 μl of 3X Laemmli buffer. Western blot samples were stored at −20°C and boiled at 95°C for 5-10 minutes before loading.

### Primary prion infection

200,000 cells were plated into 6-well plates and incubated at 37°C overnight, where each infection experiment was performed at least in duplicate. The following day, the media was replaced with 900 μl serum-free media. 100 μl of 10% prion-infected mouse brain homogenate was added for 5 hours, then 1 ml of complete media was added. This date was considered passage “0” (P0). The next day, every well was washed with 2 ml 10x PBS then 3 ml of complete media was added. Cells were grown to confluency and passaged with 0.5 ml trypsin into two 10-cm cell culture plates for P1. Cells were subsequently grown to confluency and passaged for 6-8 passages.

### Cell lysis and PK digestion for infection experiments

For infection experiments, even passages (P2, P4, P6, P8) were analyzed using Western blot. For each infected cell line replicate, one confluent plate was lysed with 1 ml 1X cell lysis buffer after washing with PBS. The plate was kept for 10 minutes, and lysate was added to a 1.5 ml microcentrifuge tube and centrifuged at 15,000g for 1 minute. 500 μL of supernatant was taken for -PK samples, where 20 μL 50X Pefabloc (Roche, # 11429868001) was added, and the samples were stored in 3 mL methanol at −20°C overnight. The remaining 500 μl of lysate, used for +PK samples, was digested with 20 μg/ml PK for 30 minutes at 37°C. After incubation, 20 μl of 50X Pefabloc was added, and the sample was stored in 3 ml of methanol overnight. The next day, samples were spun at 3,500 rpm for 25 minutes, and the pellets were resuspended in TNE buffer and 3X Laemmli buffer. -PK samples were resuspended with 100 μl TNE, and 50 μl 3X Laemmli buffer, while +PK samples were resuspended with 50 μl TNE, and 25 μl 3X Laemmli buffer. All samples were boiled for 5-10 minutes before loading.

### Immunoblot

Uninfected cell lysates being tested for Hsp110 expression were run on 10.5% acrylamide gels. Loading volume varied from 20-30 μl, where the Precision Plus Protein Dual Color Standard was used as a protein ladder. Gels were run with 1X running buffer (250 mM Tris-HCl, 2.5 M glycine, 1% (w/v) SDS) at 100 V for approximately 30 minutes (until samples passed through the stacking gel), then at 180 V for 90-120 minutes (until the sample reached the bottom of the separating gels). Gels were transferred using the Trans-Blot SD Semi-Dry Transfer Cell onto methanol-activated Amersham Hybond P 0.45 μm polyvinylene difluoride (PVDF) membranes for 60-90 minutes at 25 V. Membranes were blocked in blocking buffer for 30 minutes, after which they were incubated at 4°C overnight with relevant primary antibodies (Hsp105 - Santa Cruz ab B7 (sc-74550) at 1:250, Apg-1 – Santa Cruz mAb C6 (sc-137027) and Santa Cruz mAb B7 (sc-137007) at 1:250, Apg-2 – Santa Cruz mAb A7 at 1:250).

Infected cell lysates were run on 12.5% acrylamide gels using a similar protocol. The loading volume for -PK samples was 10 μl, and the loading volume for +PK samples was 20 μl. Brain homogenate samples were loaded with volumes of 5 μl and 10 μl for -PK and +PK, respectively. For infection experiments, 10 μl of -PK uninfected N2a cell lysate was loaded as a control. Gels were run at 100 V for 30 minutes, then at 200 V for 90 minutes, then transferred for 1 hour at 25 V. Samples were blocked in blocking buffer for 30 minutes, then incubated at 4°C overnight with 4H11 (1:500) and AC-15 (1:50,000). All membranes were visualized the next day after incubation with peroxidase-conjugated secondary antibody (goat anti-mouse at 1:5000) (Jackson Immunoresearch, #115-005-003) for 1 hour with rocking. Membranes were washed with TBS-T (100 mM Tris-HCl pH 8.0, 100 mM NaCl, 0.5% (w/v) Tween 20) for 30-40 minutes, washing for 5-minute intervals, then incubated with ECL Ultra Western HRP Substrate (MilliporeSigma, #WBULS0100) for 5 minutes. Membranes were dried and visualized using autoradiography films.

### RT-QuIC

Mouse recombinant prion protein was prepared as a substrate for RT-QuIC. Mature mouse PrP was cloned into pET41a expression vectors (EMD Biosciences) and expressed in *E. coli* Rosetta using the Novagen Overnight Express Autoinduction System 1 (MilliporeSigma, #713003). Inclusion bodies were prepared using the Novagen Bug Buster reagent (MilliporeSigma, #714563) and solubilized in 1x lysis buffer for 50 min at 23°C and then centrifuged at 16,000×g for 5 min at 23°C. Binding, refolding, and elution using an AKTA Explorer system have been described previously [33].

RT-QuIC was performed as previously described [31, 34]. 96-well plates were prepared with 98 μl of reaction mix (20mM sodium phosphate pH 6.9, 300 mM NaCl, 1 mM EDTA, 10 uM Thioflavin-T and 0.1 mg/ml recombinant mouse PrP substrate). Quadruplicate reactions were prepared with 2 μl of cell lysate samples, where each sample was diluted serially 10-fold in seed dilution buffer (0.05% (w/v) SDS in 1xPBS). The plate was sealed with Nunc Amplification Tape (Nalge Nunc International) and placed in a BMG Labtech FLUOstar Omega fluorescence plate reader that was pre-heated to 42 °C for a total of 50 h with cycles of 1-min double orbital shaking (700 rpm) and 1-min resting throughout the assay time. The Thioflavin T fluorescence signals of each well were read and documented every 15 minutes for 200 cycles, then the values of the relative fluorescence units (RFU) were plotted as the average of octuplicate reactions. The cut-off was calculated based on the average fluorescence values on the negative controls + 5 times the standard deviation.

### Filter Aided-Sample Preparation (FASP) and Mass Spectrometry (MS)

The Southern Alberta Mass Spectrometry (SAMS) FASP protocol was used for sample preparation for MS. 2-200 µg of precipitated protein was washed with acetone 3 times and centrifuged at 14,000 g for 15 min at 4°C to remove acetone. Pellets were dried and resuspended in 230 μl of Urea/Tris mix (Urea 8 M in 100 mM Tris-HCl pH 8) and vortexed for 20 minutes. 10 mM of dithiothreitol (2.3 μl of 1 M stock; Sigma-Aldrich, #D0632-5g) was added to the fully resuspended pellets and incubated at 37°C for 30 minutes, then centrifuged at 10,000 g for 5 minutes. Reduced proteins were centrifuged in Microcon YM-30 (Millipore, #MRCF0R030) at 14,000 g for 20 minutes. 200 μl of Urea-Tris was added, and the mix was centrifuged at 14,000 g for 20 minutes, then the flow-through was discarded. 100 μl of iodoacetamide (Sigma-Aldrich, #I6125-5g) 50 mM prepared in Urea-Tris was added and incubated in the dark at room temperature for 20 minutes. After the incubation, the mixture was centrifuged at 14,000 g for 20 minutes and the flow-through was discarded. 100 μl of 50 mM ammonium bicarbonate was added and centrifuged at 1,400 g for 15 minutes, and this step was repeated twice. 40 μl of trypsin (20 ug trypsin resuspended in 10 μl of 10 mM acetic acid resuspension buffer, in 80 μl of 50 mM ammonium bicarbonate) was added and centrifuged in the YM-30. Samples were covered with aluminum foil and incubated overnight at 37°C. The following day, samples were centrifuged at 100 g for 5 minutes then left at room temperature for 10 minutes. The YM-30 membranes were transferred to a new collection tube, and samples were centrifuged at 14,000 g for 10 minutes. 40 μl of 50 mM sodium bicarbonate was added and samples were centrifuged at 14,000 g for 10 minutes, repeating twice.

Peptides were analyzed on an Orbitrap Fusion Lumos Tribrid mass spectrometer (Thermo Scientific) operated with Xcalibur (version 4.4.16.14) and coupled to a Thermo Scientific Easy-nLC (nanoflow Liquid Chromatography) 1200 system. Peptides were loaded onto a C18 trap (75 um x 2 cm; Acclaim PepMap 100, P/N 164946; ThermoScientific) at a flow rate of 2 μl /min of solvent A (0.1% formic acid in LC-MS grade water). Peptides were eluted using a 120 min gradient from 5 to 40% (5% to 28% in 105 min followed by an increase to 40% B in 15 min) of solvent B (0.1% formic acid in 80% LC-MS grade acetonitrile) at a flow rate of 0.25 μl/min and separated on a C18 analytical column (75 µm x 50 cm; PepMap RSLC C18; P/N ES803; ThermoScientific). Peptides were then electrosprayed using 2.1 kV voltage into the ion transfer tube (300°C) of the Orbitrap Lumos operating in positive mode. The Orbitrap first performed a full MS scan at a resolution of 120000 FWHM to detect the precursor ion having a m/z between 375 and 1575 and a +2 to +7 charge. The Orbitrap AGC (Auto Gain Control) and the maximum injection time were set at 4e5 and 50 ms, respectively. The Orbitrap was operated using the top speed mode with a 3 sec cycle time for precursor selection. The most intense precursor ions presenting a peptidic isotopic profile and having an intensity threshold of at least 5,000 were isolated using the quadrupole and fragmented with HCD (30% collision energy) in the ion routing multipole. The fragment ions (MS2) were analyzed in the ion trap at a rapid scan rate. The AGC and the maximum injection time were set at 1e4 and 35 ms, respectively, for the ion trap. Dynamic exclusion was enabled for 45 sec to avoid the acquisition of the same precursor ion having a similar m/z (plus or minus 10 ppm).

## Results

### Transient knockdown of Hsp110 family members reduces PrP^Sc^ in persistently prion-infected cell lines

To test whether manipulation of Hsp110 family members affects prion infection in persistently infected cells, we transiently knocked down Apg-1 and Hsp105 in mouse neuroblastoma (N2a) cells infected with 22L prions (22L-ScN2a) (**Fig. 1A**). Cells were co-transfected with siRNA against Apg-1 and Hsp105 (KD) or transfected with non-targeting siRNA (CT). Transfections were performed twice at 48-hour intervals, and cell lysates were collected 96 hours after the first transfection. Lysates were subjected to proteinase K digestion or not (+/- PK), followed by immunoblot analysis to compare PrP^Sc^ levels in control and knock-down cells. Our results showed decreased PrP^Sc^ levels in knock-down cells following transient co-transfection of siRNA targeting Hsp105 and Apg-1. To validate this observation in other cell lines, we repeated the experiment with CAD5 cells persistently infected with RML prions (RML-ScCAD5) (**Fig. 1B**), and mouse embryonic fibroblasts infected with 22L prions (22L-ScMEF) (**Fig. 1C**). Our analysis revealed a reduction of PrP^Sc^ levels in each cell line following the transient knockdown. Densitometric analysis validated these observations, with PrP^Sc^ levels reduced by 48% in 22L-ScN2a cells (*p* = 0.040; KD *vs* CT cells; mean ± SD, arbitrary units: 0.1628 ± 0.0322 vs 0.3134 ± 0.0809; *n* = 3/group) (**Fig. 1D**). Reductions of similar magnitude were observed in RML-ScCAD5 cells (49%; *p* = 0.027; *n* = 3/group) and 22L-ScMEF cells (54%; *p* = 0.037; *n* = 4/group) **(Fig 1D)**. Densitometric analysis also confirmed the reduced expression of Hsp105 in each cell line **(Fig 1E)**. 22L-ScN2a cells demonstrated a 77% reduction (*p* < 0.001; *n* = 3/group), while RML-ScCAD5 cells and 22L-ScMEF cells demonstrated 67% and 46% reductions, respectively (*p* = 0.013; *n* = 3/group (RML-ScCAD5); *p* = 0.045; *n* = 4/group (22L-ScMEF)) **(Fig 1E)**.

**Fig 1.**
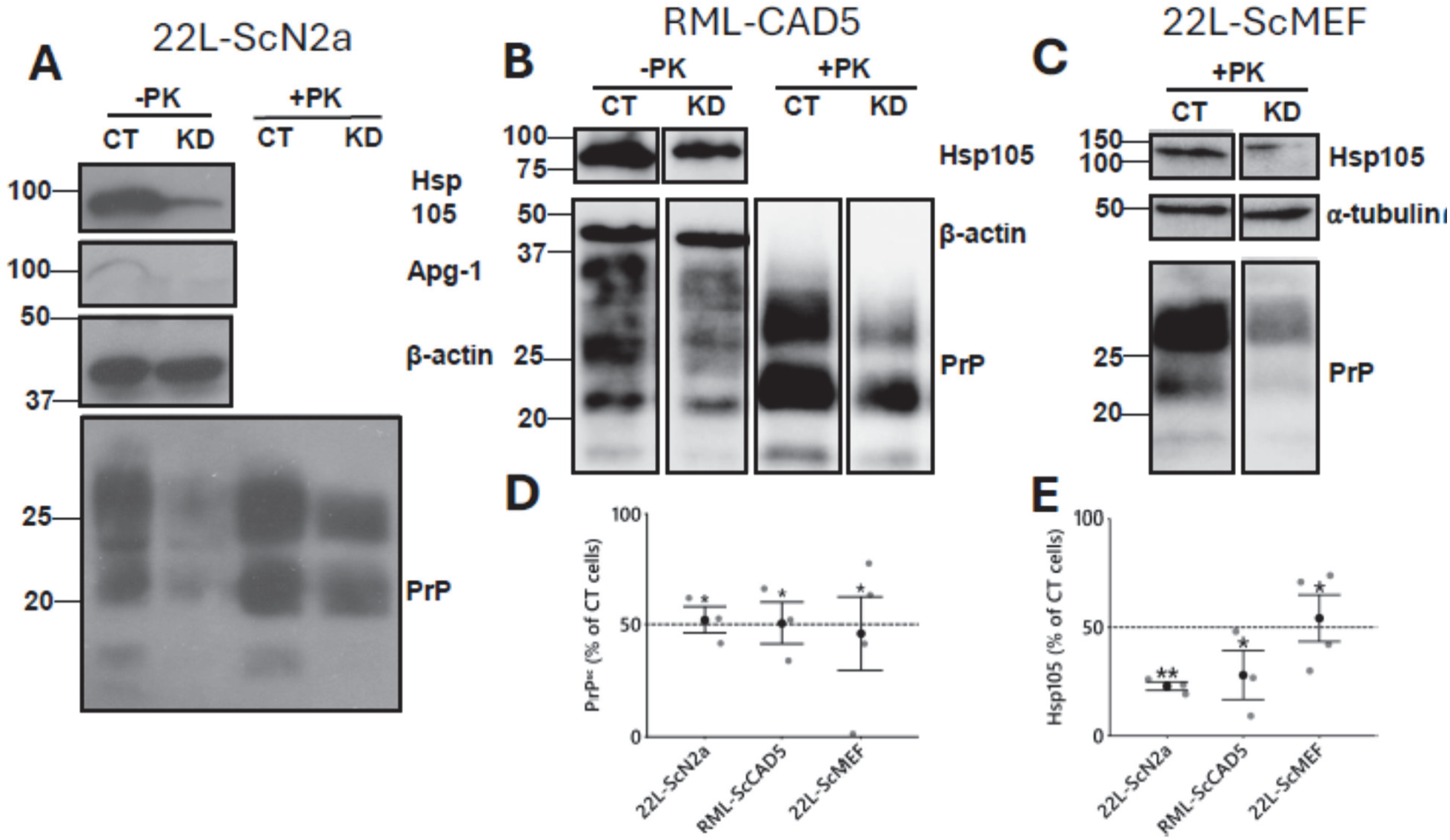
Reduced PrP^Sc^ levels in various prion-infected cell lines following transient knockdown of Apg-1 and Hsp105. (A-C) Representative immunoblots illustrating PrP^Sc^ reduction in **(A)** 22L-ScN2a, **(B)** RML-ScCAD5, and **(C)** 22L-ScMEF cells 96 hours after Hsp110 knockdown (KD). Persistently prion-infected cells were co-transfected with siRNA against Apg-1 and Hsp105 (KD), or with non-targeting siRNA (CT). Transfections were carried out twice – 48 hours apart – and cells were collected 96 hours after the first transfection. Post-nuclear cell lysates were analyzed via immunoblot, and PK digestion was used to distinguish PrP^Sc^ from total PrP. PK-resistant PrP^Sc^ (+PK) shows up as un-glycosylated (19 kDa), mono-(23/24 kDa) and di-glycosylated bands (27-30 kDa). Beta-actin (β-actin) or alpha-tubulin (α-tubulin) were used as loading controls. **(D-E)** Densitometric quantification (mean ± SEM) demonstrates a significant reduction in **(D)** PrP^Sc^ and **(E)** Hsp105 in all three KD cell lines compared to WT. Data were normalized to loading controls and shown as a percentage of the CT cells. Statistical analysis was performed using a Student’s *t-test* (* *p*<0.05; *** *p*<0.001).

Overall, these results show that knockdown of Hsp110 family members reduces PrP^Sc^ levels in three different cell models, persistently infected with two types of mouse-adapted scrapie prions.

We next examined the effect of Hsp110 overexpression on pre-existing prion infection. N2a cells persistently infected with 22L prions (22L-ScN2a) were transfected with an empty vector (CT) or co-transfected with expression vectors encoding Apg-1 and Hsp105. Transfections were carried out twice, and cells were collected 96 h after the first transfection. Lysates were subjected to PK digestion or not (+/-PK), followed by immunoblot analysis (**Fig. S1**). Densitometric analysis revealed a twofold increase in PrP^Sc^ levels in co-transfected cells compared to control-treated cells (p = 0.037; n = 5 (CT), n = 6 (OE)). Next, we transfected 22L-ScN2a cells with 1 μg of a mixture of expression vectors encoding all three Hsp110 family members, or with 2 μg of said mixture. Densitometric analysis (**Fig. S2**) revealed a dose-dependent increase in PrP^Sc^ levels, which was statistically significant for cells transfected with 2 μg of expression vectors (*p* = 0.040; *n* = 3/group). These data demonstrate that transient overexpression of Hsp110 family members increases PrP^Sc^ in stably prion-infected cells, having an effect opposite to that of transient knockdown.

Taken together, manipulation of Hsp110 in persistently infected cells affects the levels of PrP^Sc^.

### Generation of cell lines with stable knockout of Hsp110 members

Having observed that Hsp110 expression affects PrP^Sc^ levels in persistently infected cells, we next wanted to see whether Hsp110 is also required for *de novo* prion infection. Thus, we generated N2a cell lines devoid of either Hsp105, Apg-1, or Apg-2 using a CRISPR-Cas9 system. To generate an Apg-1 knock-out (KO) cell line, uninfected N2a cells (WT) were treated with guide RNAs (gRNAs) targeting three loci on exon 1 of the *Hspa4l* gene **(Fig 2A)**. Multiple single-cell clones were isolated following treatment by limiting dilution; however, initial protein analysis revealed incomplete knockouts in each of the isolated clones (*data not shown*). Therefore, CRISPR treatment was repeated on isolated clones, aiming to knock out the protein completely. Single-cell cloning was not performed following the re-treatment of the isolated clones. As illustrated in **Fig. 2B**, immunoblot analysis revealed that Apg-1 was not detectable in the re-treated cells. Apg-1 deletion was also confirmed via DNA sequencing analysis. To confirm KO across all alleles in these aneuploid cells, PCR products were subcloned and at least 20 plasmid clones analyzed. Four different patterns of deletions and insertions were observed, all resulting in frameshifts and premature stop codons (**Fig. 2A**). Finally, mass spectrometry was used to validate the complete knockout of Apg-1, indicating also that Apg-1 was not significantly up- or down-regulated in other Hsp110 knockouts **(Fig. 2C).**

**Fig 2.**
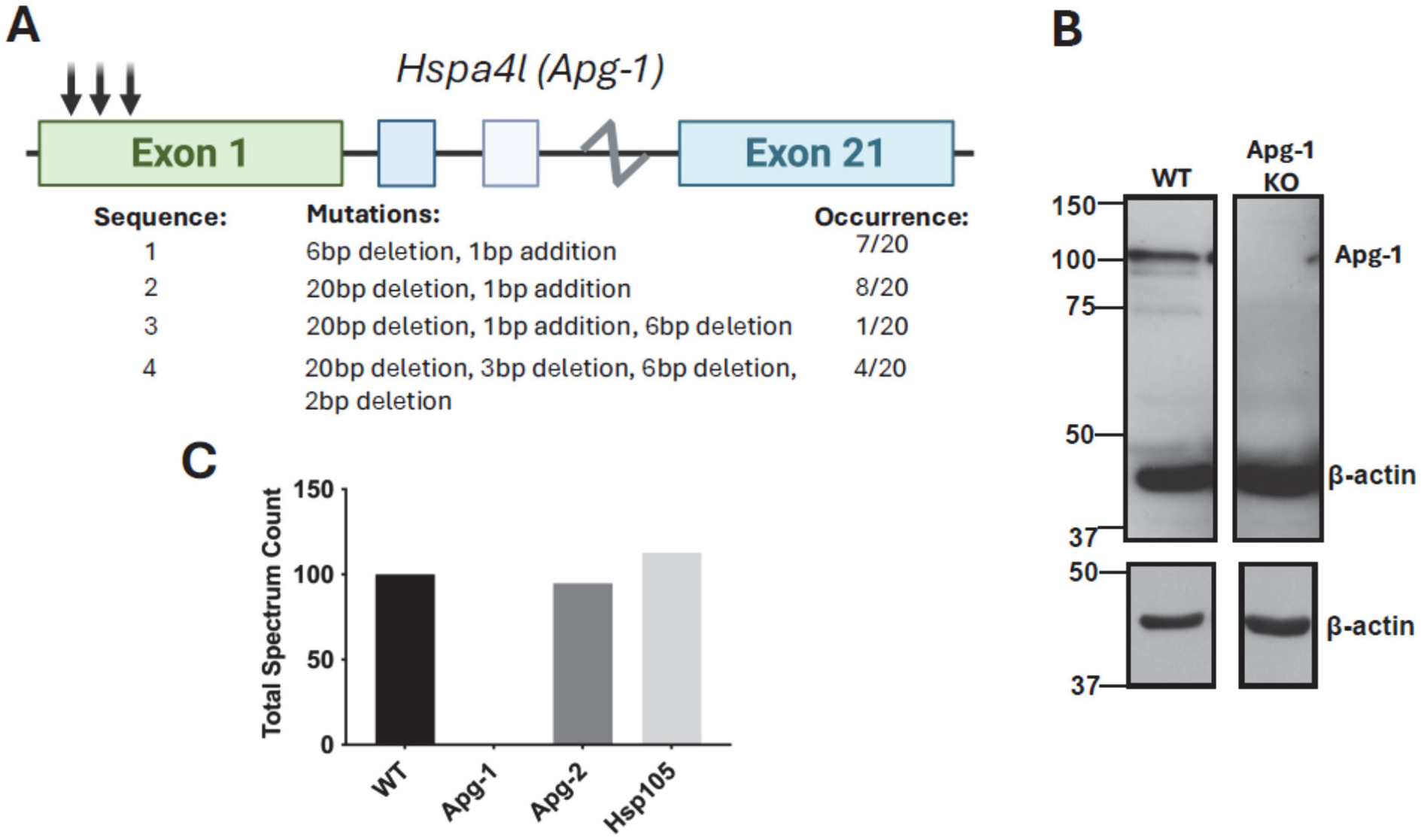
Generation of Apg-1 knockout N2a cells using CRISPR/Cas9. **(A)** Schematic illustrating gRNA target sites on the *Hspa4l* gene encoding Apg-1. The induced mutation patterns are described, along with the frequency observed in the 20 plasmid clones analyzed. **(B)** Immunoblot confirming the absence of Apg-1 in CRISPR/Cas9 treated cells. Blots were probed for Apg-1 using mAb B7. Wild-type N2a (WT) cell lysate and beta-actin (β-actin) were included as positive and loading controls, respectively. The lower blots show a shorter exposition for the actin signal. **(C)** Bar graph illustrating consistent Apg-1 levels across WT, Apg-2 KO, and Hsp105 KO cells. As expected, Apg-1 KO cells are devoid of Apg-1 protein. Total spectrum count was derived following mass spectrometry and serves as a proxy measurement for protein levels. Visualized in GraphPad Prism 11.

To generate an Apg-2 KO cell line, uninfected N2a cells were treated with gRNA’s targeting three loci on exon 1 of the *Hspa4* gene **(Fig. 3A)**. Several single-cell clones were isolated following treatment, and one clone was chosen for subsequent experiments and subjected to further immunoblot and DNA sequencing analysis. Immunoblot analysis demonstrated the lack of Apg-2 (**Fig. 3B**). Complete Apg-2 deletion was confirmed via DNA sequencing analysis of 22 clones, detecting 5 insertion-deletion patterns all resulting in frameshift and premature stop codons **(Fig 3A).** Mass spectrometry confirmed that the Apg-2 KO was complete and that this KO had no effects on the remaining Hsp110 proteins **(Fig 3C).** To generate a Hsp105 KO cell line, N2a cells were treated with gRNAs targeting one locus on exon 1 and two loci on exon 7 of the *Hsph1* gene **(Fig. 4A)**. Single-cell clones were isolated and analyzed via immunoblot to examine Hsp105 expression. The majority of the isolated clones demonstrated a complete lack of Hsp105 (*data not shown*). The single-cell clone chosen for subsequent experiments underwent further immunoblot and DNA sequencing analysis (**Fig. 4A** and **B**). Sequencing of the plasmid library indicated 2 gene editing patterns in the 15 analyzed DNA clones **(Fig. 4A)**. Both analyses confirmed the knockout of Hsp105 in the chosen single-cell clone (**Fig. 4A** and **B**). Unfortunately, Hsp105 did not show up in the MS analysis.

**Fig 3.**
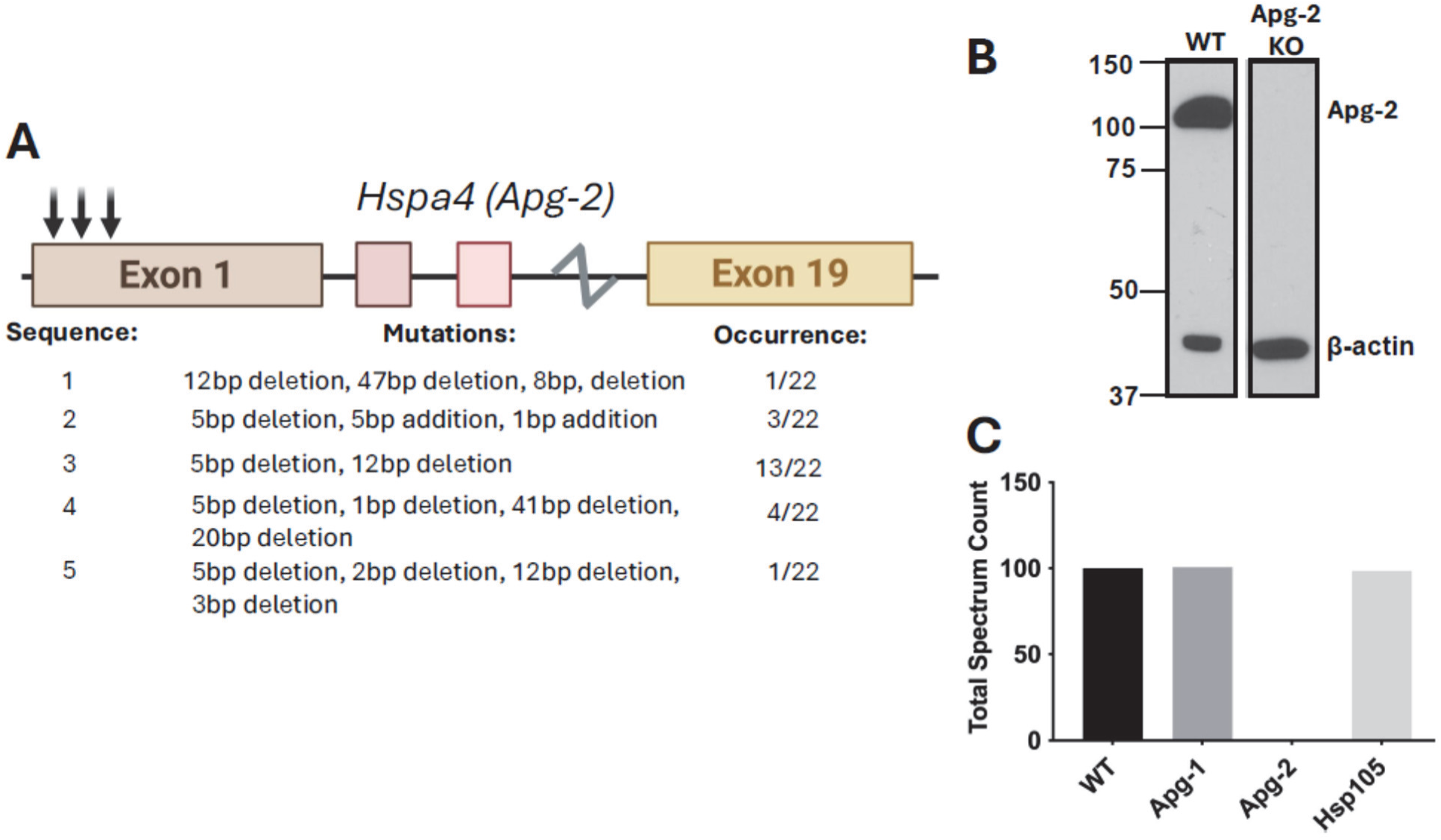
Generation of Apg-2 knockout N2a cells using CRISPR/Cas9. **(A)** Schematic illustrating gRNA target sites on the *Hspa4* gene encoding Apg-2. The induced mutations are described, along with the frequency of occurrence in analyzed plasmid clones. **(B)** Immunoblot confirming the absence of Apg-2 in CRISPR/Cas9 treated cells. The blot was probed for Apg-2 using mAb A7. Wild-type N2a (WT) cell lysate and beta-actin (β-actin) were included as positive and loading controls, respectively. **(C)** Bar graph illustrating consistent Apg-2 levels across WT, Apg-1 KO, and Hsp105 KO cells. As expected, Apg-2 KO cells are devoid of Apg-2 protein. Total spectrum count was derived following mass spectrometry and serves as a proxy measurement for protein levels. Visualized in GraphPad Prism 11.

**Fig 4.**
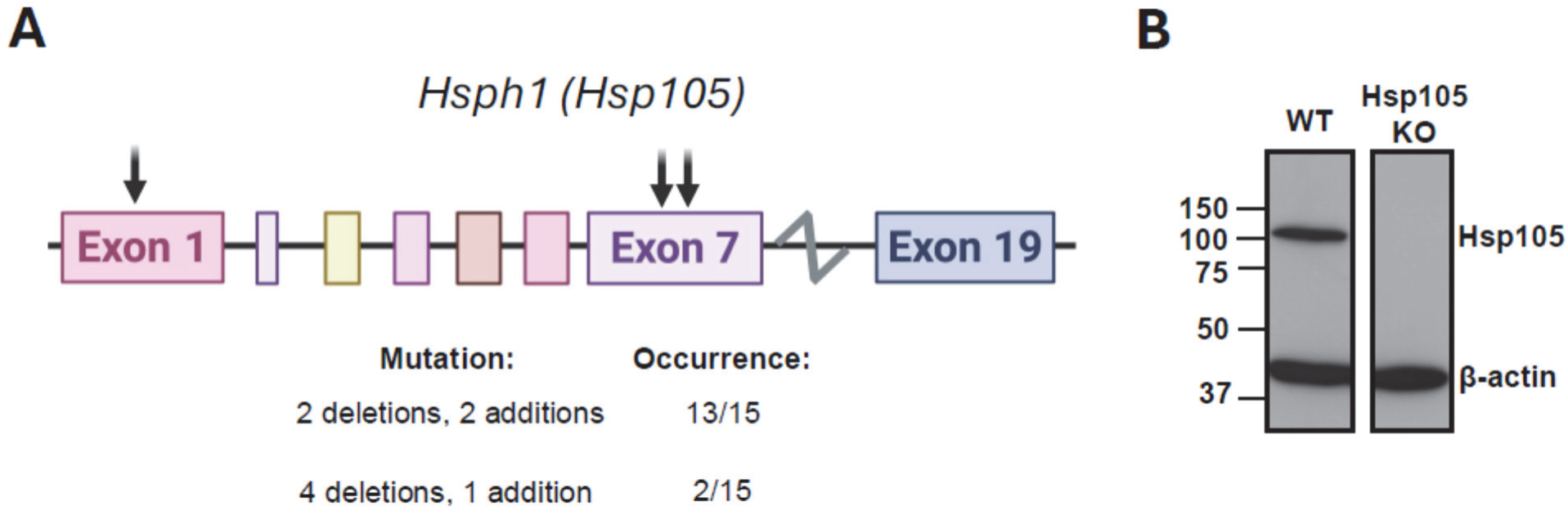
Generation of Hsp105 knockout N2a cells using CRISPR/Cas9. **(A)** Schematic illustrating gRNA target sites on the *Hsph1* gene encoding Hsp105. The induced mutations are described, along with the frequency of occurrence. **(B)** Immunoblot confirming the absence of Hsp105 in CRISPR/Cas9 treated cells. Blot was probed for Hsp105 using mAb B7. Wild-type N2a (WT) post-nuclear lysate and beta-actin (β-actin) were included as positive and loading controls, respectively.

Next, we wanted to ensure that the genetic knockouts did not have deleterious effects on cellular viability. Therefore, we performed LDH and XTT assays to investigate cell viability in each of the Hsp110 KO cell lines. The LDH assay measures cytotoxicity, while the XTT assay permits the quantification of cell viability, proliferation, and cytotoxicity more broadly. For the LDH assay, cell culture media were collected from each Hsp110 KO and tested. For the XTT assay, reagents were added directly to cultured cells. There were no statistically significant differences observed in the LDH assay between KO and WT cells **(Fig. 5A).** The XTT assay revealed no statistically significant differences between the Hsp105 and Apg-1 KO cells in comparison to WT cells **(Fig 5B)**. However, there was a statistically significant increase in cell viability in the Apg-2 KO cells **(Fig 5B)**.

**Fig 5.**
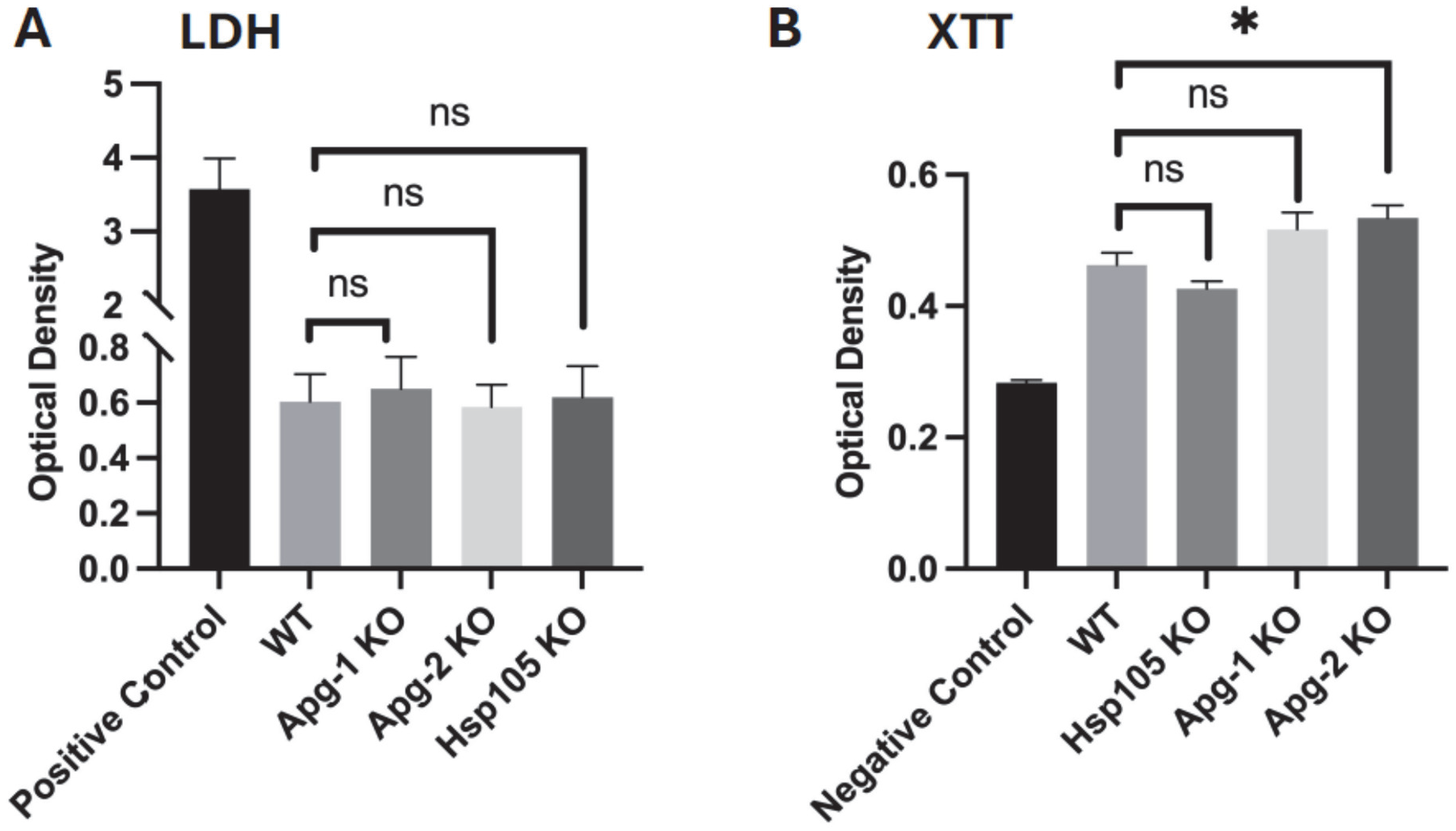
Cell viability is not negatively affected by Hsp110 KO. **(A)** Bar graph (mean ± SEM) depicting cytotoxicity measured by LDH release across WT and Hsp110 knockout cells. Cell culture medium was collected, and LDH levels were elucidated via colorimetric analysis. Optical density measurements are representative of LDH levels within the sampled medium. The positive control represents sampled medium from cells treated with Triton X-100 **(B)** Bar graph (mean ± SEM) depicting the cell viability measured by XTT assay of WT and Hsp110 knockout cell lines. Cells were cultured with the XTT assay reagents, and the amount of XTT reduction was quantified via colorimetric analysis. Optical density measurements are representative of the amount of XTT reduction within the cultured cells. A slight but significant increase was observed in the Apg-2 KO cell line. The negative control represents cells treated with Triton X-100. Both graphs were visualized in GraphPad Prism 11. Statistical analyses were performed using a *One-way ANOVA* with a Dunnett’s test for pair-wise comparisons (“ns” signifies non-significance; * p<0.05).

In conclusion, we successfully generated Apg-1, Apg-2 and Hsp105 KO cell lines. The genetic modifications did not have deleterious effects on cellular viability.

### Hsp110 KOs reduce levels of PrP^Sc^ in *de novo* prion infection differently

Having established that KOs do not affect cell viability, we next examined the effect of Hsp110 KO on *de novo* prion infection. Cells were infected with brain homogenates from terminally sick mice infected with 22L prions. Cells were passaged 6-8 times, with one passage roughly corresponding to 5 days, and alternating passages were lysed and analyzed with immunoblot and RT-QuIC to assess PrP^Sc^ levels and seeding activity, respectively.

Infection experiments were done in at least 3 bio-replicates. A representative experiment is shown in **Fig. 6**, depicting WT cells and all 3 KOs over passages 2 to 8. In this experiment at passage 2, a PrP^Sc^ signal was detectable only in WT and Apg-2 KO cells (**Fig. 6A**). PrP^Sc^ signals showed up in all tested passages for WT and Apg-2 KO cells, and passages 4 to 8 for Hsp105 KO cells (**Fig. 6A-D**). For Apg-1 KO cells, PrP^Sc^ was only detectable in passage 6 in this experiment, and at lower levels compared to WT and Apg-2 KO cells (**Fig. 6C**). These results reveal distinct patterns of infection in each of the Hsp110 KOs across various passages. Densitometric analysis of PrP^Sc^ signals of all bio-replicates compared to PrP^Sc^ levels in WT control cells validated these findings (**Fig. 7**). The Apg-1 KO demonstrated robust reduction of PrP^Sc^ levels at each passage (**Fig 7A)**, with early passage representing passage 2, middle passage 4, and late passage 6 and/or 8 **(Fig 7A)**. Interestingly, an almost total loss of PrP^Sc^ signal is observed at late passages **(Fig 7A).** In contrast, the Apg-2 KO only demonstrates lower PrP^Sc^ levels at early passages **(Fig 7B).** At the middle and later passages, PrP^Sc^ levels were comparable to those of WT cells **(Fig 7B).** The Hsp105 KO demonstrated reduced PrP^Sc^ levels at each passage. More pronounced reductions were visible in the early and middle passages, suggesting a slower onset of infection **(Fig 7C).** Overall, immunoblot results indicate that KOs of Apg-1 and Hsp105 have strong and robust effects on *de novo* prion infection, detectable even after 8 passages. Knock-out of Apg-2 was much less effective and only detectable at early passages.

**Fig 6.**
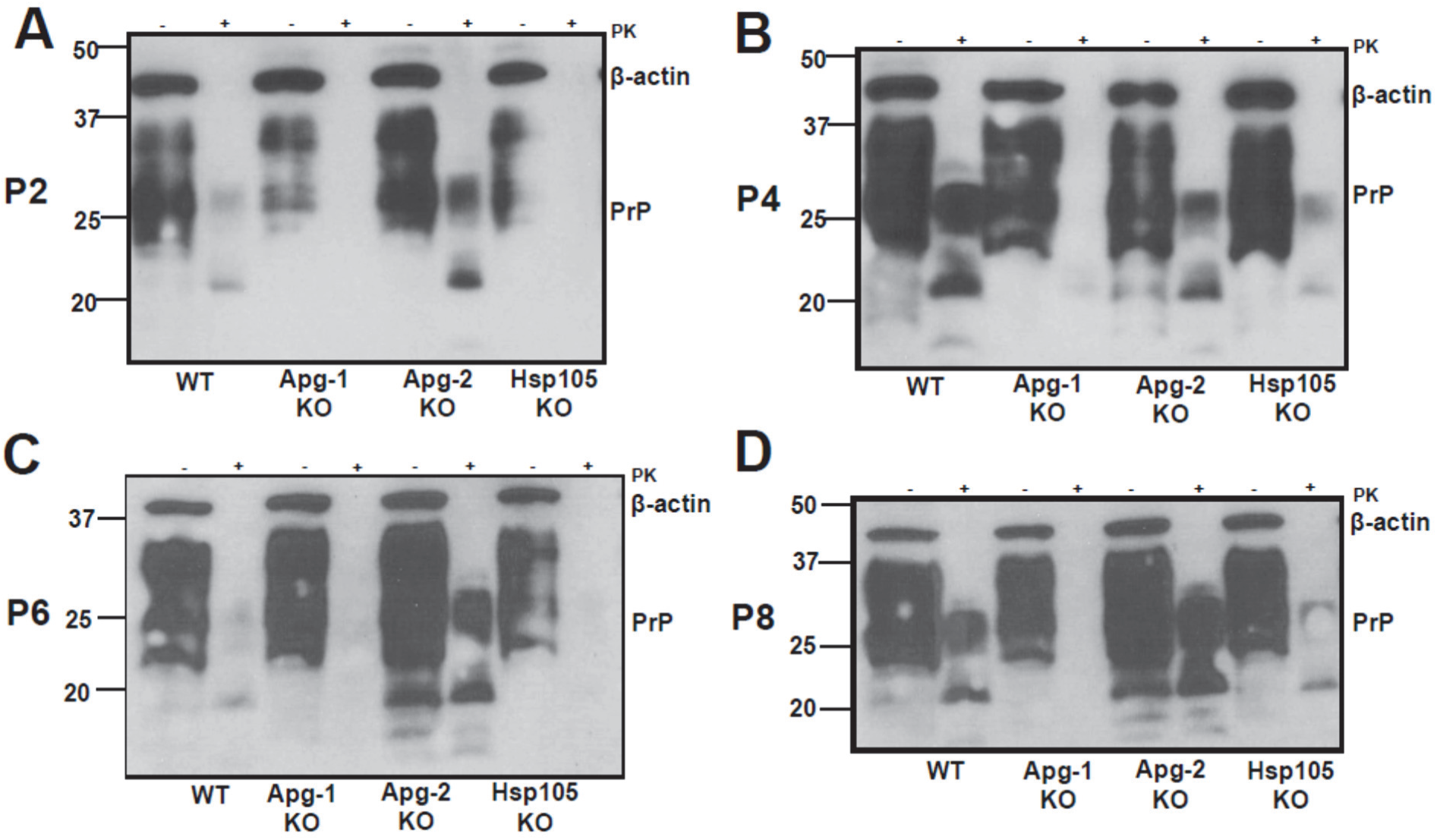
Representative immunoblots demonstrating effects of Hsp110 KO on primary prion infection. (A-D) PrP^Sc^ signals in WT and all three Hsp110 KO cell lines were measured at P2 **(A)**, P4 **(B)**, P6 **(C)**, and P8 **(D).** Cells were exposed to 22L-infected mouse brain homogenate and passaged every 5 days. Cell lysates were collected at the indicated passages, samples were treated with or without PK (+/-PK), separated on SDS-PAGE and analyzed by immunobloting. mAb 4H11 was used for PrP identification. PK-resistant PrP^Sc^ (+PK) shows up as un-glycosylated (19 kDa), mono-(23/24 kDa) and di-glycosylated bands (27-30 kDa). Beta-actin (β-actin) was included as a loading control, co-probed in the same immunoblot.

**Fig 7.**
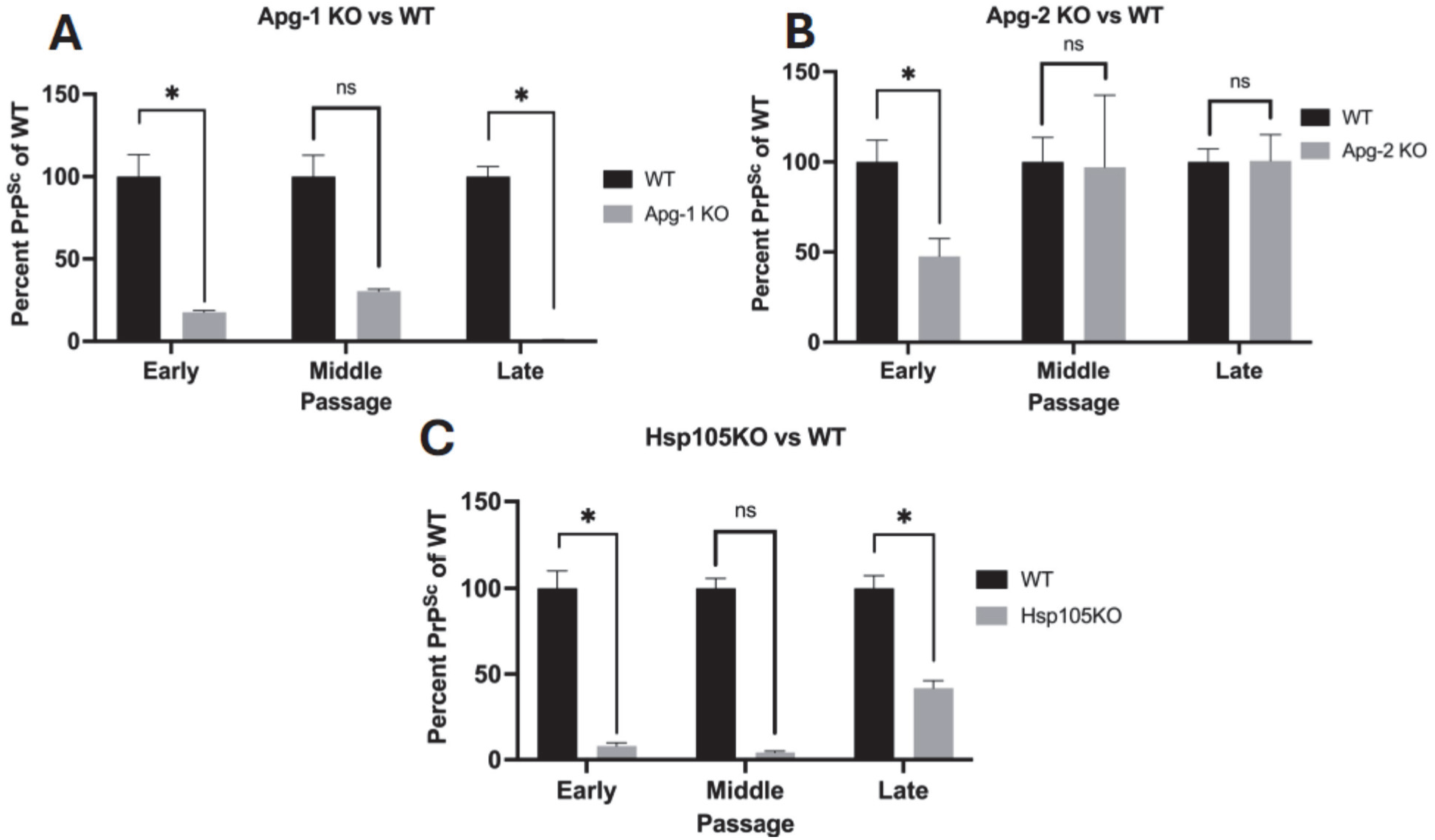
Densitometric analysis elucidating distinctive patterns of prion infection in each Hsp110 KO cell line. (A-C) Bar graphs (mean ± SEM) illustrating changes in PrP^Sc^ levels measured by immunoblot across early (P2-P3), middle (P4), and late passages (P6-P8) in Hsp105 KO **(A)**, Apg-2 KO **(B)**, and Apg-1 KO **(C)** cell lines. PrP^Sc^ signal intensity was measured using *ImageJ* and normalized to WT cells for the various passages. At least 3 independent bio-replicates were analyzed, often with 2 technical replicates. PrP^Sc^ bands were normalized to beta-actin before densitometric analysis. Statistical analyses were performed using a one-tailed paired *t-test* (“ns” signifies non-significance; * p<0.05). All graphs were visualized in GraphPad Prism 11.

To correlate immunoblot findings to prion seeding activities, we performed RT-QuIC assays. At passage 2, time to cross the cut-off was shortest for Apg-2 KO cells, and longest for Apg-1 KO cells, indicating lowest seeding activity in Apg-1 KO cells and replicating closely the immunoblot pattern in Fig. 6 (**Fig. 8A**). Quantification of maximum RFU (**Fig. S3A**) and area under the curve (**Fig. S4A**) values validated this finding. This pattern of Apg-1 KO cells having the lowest seeding activity was also consistent over passages 4 to 8, with less pronounced differences to WT cells at passage 8 (**Fig. 8B-D**; **Fig. S3B-D**, **Fig. S4B-D**). For passage 8, there could be a dissociation of PK-resistant PrP^Sc^ and prion seeding activity patterns in Apg-1 KO cells. An inverse prion seeding activity pattern is observed for Apg-2 KO cells with signals almost stronger than WT cells over all 4 tested passages, replicating PrP^Sc^ signals in immunoblot analysis **(Fig 8A-D, Fig. S3A-D, Fig. S4A-D).** The Hsp105 KO cells demonstrate only slightly reduced seeding activity relative to WT cells in most passages **(Fig 8, Fig. S3** and **S4).**

**Fig 8.**
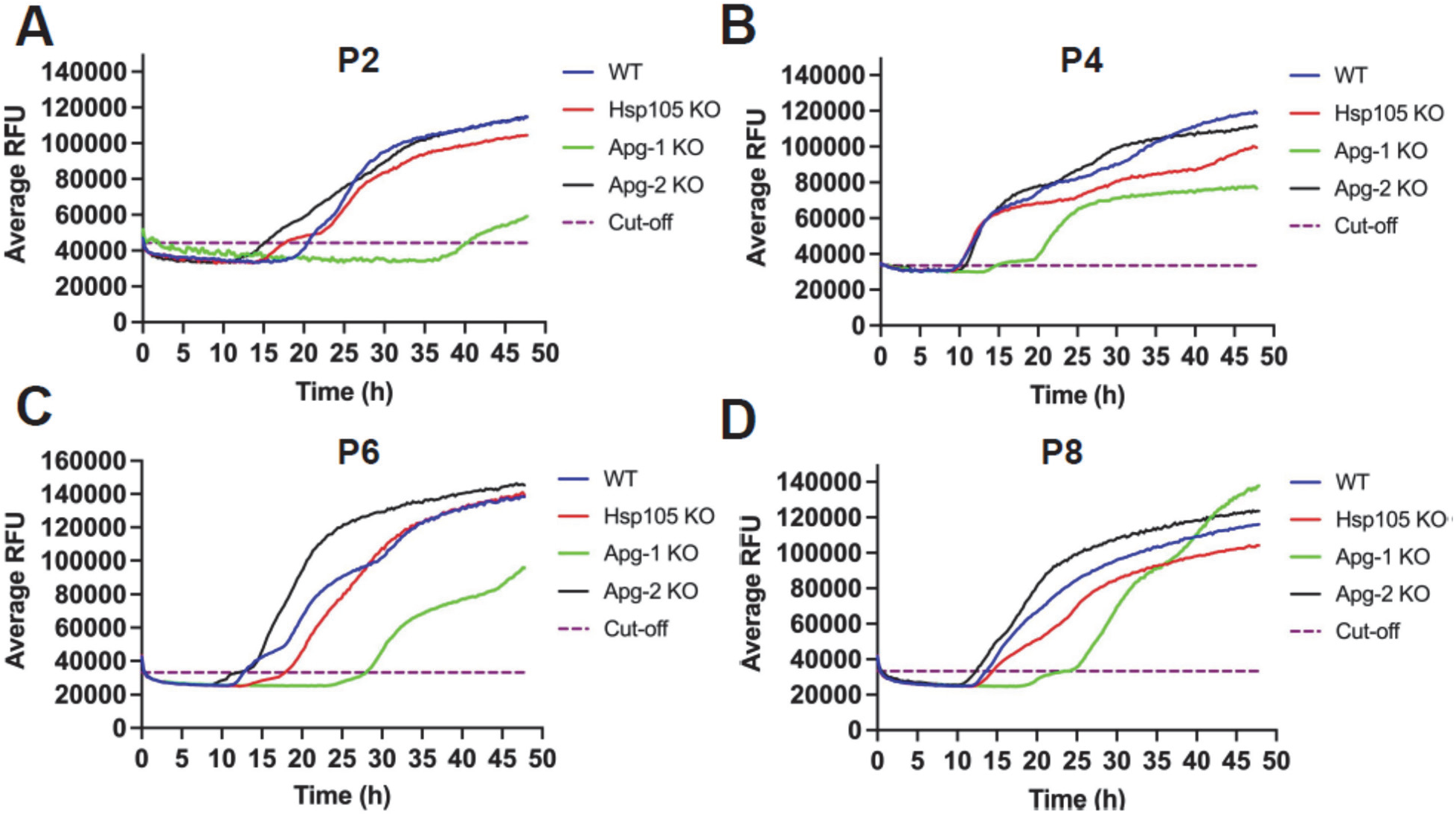
Prion seeding activity in Hsp110 KO cell lines compared to WT cells. (A-D) Kinetic curves illustrate prion seeding activity in WT and Hsp110 KO cell lines at P2 **(A)**, P4 **(B)**, P6 **(C)**, and P8 **(D)**. Each cell line was exposed to 22L-infected mouse brain homogenate and was subsequently passaged every 5 days. Cell lysate from the mentioned passages was collected and utilized for RT-QuIC analysis (1:20 dilution shown). Mouse recombinant PrP was used as a substrate. The average Thioflavin-T fluorescence is plotted as a function of time. Average relative fluorescence units (RFU) are plotted on the *y*-axis, and time (h) is depicted on the *x*-axis. The assay was performed in quadruplicates and was visualized in GraphPad Prism 11. Samples were considered positive when 2 out of 4 wells crossed the threshold, defined as the average relative fluorescent unit (RFU) of the negative control group plus five times its standard deviation.

Taken together, knock-out of Apg-1 reduced prion seeding activity in all analyzed passages, recapitulating results from the immunoblot analysis, demonstrating that Apg-1 is a modifier of cellular prion infection.

## Discussion

The role of the Hsp70 and Hsp110 molecular chaperone families in neurodegenerative diseases remains controversial. While one body of research suggests that these chaperones help alleviate disease progression, another highlights their potential to potentiate disease pathogenesis. In support of the former are several studies characterizing the Hsc70/DNAJB1/Apg-2 disaggregase system and its beneficial effects. This system efficiently disassembles alpha-synuclein fibrils and promotes fibril conversion into non-toxic monomers [24, 25, 35, 36]. This disaggregation machinery has demonstrated substrate versatility, as it can also disentangle huntingtin fibrils *in vitro* [37, 38]. Importantly, Rampelt and colleagues demonstrated that substituting Apg-2 with either Apg-1 or Hsp105 did not alter the disaggregation machinery’s efficacy, suggesting all three homologues have a similar role within this system [39]. Furthermore, genetic knockout and overexpression studies suggest that both Apg-1 and Hsp105 help alleviate neurodegenerative disease pathology. Hsp105 KO mice exhibited an enhancement in the pathological features associated with Alzheimer’s disease [17]. Moreover, Apg-1 overexpression has been found to reduce alpha-synuclein pathology in both cellular and mouse models of Parkinson’s disease [40]. Lastly, some research suggests that the Hsp70 disaggregase system may protect against prion disease progression. Park and colleagues demonstrated that Hsp70 overexpression reduced PrP^Sc^ levels in prion-infected cells, whereas Hsp70 silencing increased them [41]. Furthermore, in a Creutzfeldt-Jakob disease Drosophila model, Hsp110 knockdown enhanced prion-induced neurotoxicity, whereas overexpression attenuated neurotoxicity and ablated seeding activity [6].

As mentioned earlier, another body of research posits that the Hsp70 and Hsp110 molecular chaperone families potentiate neurodegenerative disease pathogenesis. Our results support this research by showcasing the beneficial role of Hsp110 reduction in several cell models of prion infection. Our investigation began with the transient KD of both Apg-1 and Hsp105 in three different stably-infected prion cell lines. Upon western blot analysis, we observed significant PrP^Sc^ reductions in each cell line **(Fig 1)**. Accordingly, overexpression of Apg-1 and Hsp105 in stably-infected 22L-ScN2a cells contributed to a significant increase in PrP^Sc^ levels **(Fig S1)**. In the same cell line, overexpression of all three Hsp110s homologues led to a dose-dependent increase in PrP^Sc^ levels **(Fig S2)**. We next examined the effect of Hsp110 modulation on *de novo* prion infection using N2a cell lines devoid of either Hsp105, Apg-1, or Apg-2. Immunoblot results revealed reductions in PK-resistant PrP^Sc^ levels in each Hsp110 KO, albeit to varying degrees. The Apg-1 KO produced robust reductions across all passages. These results were recapitulated in our RT-QuIC analyses, where we observed consistently low seeding activity across various passages. Like the Apg-1 KO, substantial PrP^Sc^ reductions at each passage were also observed in the Hsp105 KO. Interestingly, more pronounced reductions were visible in the early and middle passages, suggesting a slower onset of infection.

Overall, our work utilized various cell models of prion infection and detection methods to characterize the role of mammalian Hsp110 chaperones in cellular prion infection. Our findings lend credence to the notion that these proteins may potentiate prion disease pathogenesis. This concept draws strong precedent from seminal work in yeast, where the AAA+ disaggregase Hsp104 was shown to facilitate prion propagation. Key evidence for this comes from Hsp104 depletion studies, where reduction of the disaggregase markedly impairs the proliferation of certain yeast prion strains [14, 42–44]. Furthermore, Hsp104 deletion causes considerable increases in the size of prion aggregates and polymers [45, 46]. Building from this body of work, direct evidence demonstrates that Hsp104-driven disaggregation enhances prion seeding activity to a degree that is comparable to physical sonication [47, 48]. Specific mutations that hinder the threading mechanism of Hsp104, which underlies its disaggregation ability, significantly inhibit yeast prion propagation [49].

Overall, cumulative evidence suggests that yeast prions propagate via Hsp104-mediated disaggregation of larger agglomerates into seeding-competent species [50, 51]. Several studies, including our present work, have established a similar mechanism in higher organisms, such as nematodes and mammals. In these organisms, Hsp70 and Hsp110 chaperones have been implicated in the breakdown of disease-associated fibrils into seeding-competent species. For instance, in an Alzheimer’s *C.elegans* model, Hsp110 depletion attenuated amyloid beta aggregation, whereas overexpression promoted aggregation and potentiated neurodegeneration [16]. Furthermore, Tittelmeier and colleagues found that Hsp110 knockdown reduced alpha-synuclein aggregation, toxicity, and transmission in a *C.elegans* model of Parkinson’s disease [15]. An *in vitro* study further clarified these results by demonstrating that the Hsc70/DNAJB1/Apg-2 disaggregase system could efficiently break down tau fibrils and promote the formation of small seeding-competent species [52]. In a study conducted with mammalian prions, it was observed that Hsp105 and Apg-2, in tandem with Grp78 and DNAJC1, promoted the disaggregation of 22L prions into small particles with seeding activity in acute prion infection [27].

Collectively, these findings suggest that disaggregase-mediated seed generation is a driver of prion propagation. Building on this foundation, our current work suggests that mammalian Hsp110 chaperones promote cellular prion infection. Specifically, we propose that Hsp110 knockdown or knockout impairs the mammalian disaggregase system, thereby preventing the formation of seeding-competent species that can propagate prions. While this disaggregation model offers a compelling framework, it likely provides only a partial explanation for our results, given the potential crosstalk between heat shock proteins and e.g. autophagy. Therefore, we suggest that this connection also contributes to our observed results. In support of this supposition are several studies that elucidate the potential relationship between the aforementioned proteostatic mechanisms. For instance, Dokladny and colleagues observed that deletion of heat shock transcription factor 1 (HSF1) increases basal autophagy levels [53]. Conversely, treatment with a constitutively active form of HSF1 inhibits autophagy induction in heat-treated cells [54]. In concert with these studies, Alhasan et al. observed that Hsp70 inhibition potentiates autophagy via the modulation of several autophagy-associated molecules [55]. The authors also observed that genetic or pharmacological induction of Hsp70 promoted autophagy downregulation [55]. Moreover, Hsp110 reduction potentiates autophagy, whereas overexpression has an inhibitory effect [16]. Altogether, these findings suggest that autophagy induction may play a role in our observed results, as recent work from our lab suggests that autophagy is crucial for cellular prion clearance [56]. Furthermore, our group has described various chemical compounds that induce autophagy and decrease prion infection in cell culture and mouse models [57–62].

Given the variable effects of each Hsp110 knockout on prion infection, the autonomous activities of the individual proteins warrant independent consideration. Structurally, these proteins share significant sequence identity, although key differences exist in certain regions, such as the substrate-binding domain [63, 64, 65]. These structural differences provide the foundation for the functional variations that exist between these proteins. Functional differences between the mammalian Hsp110s were demonstrated in studies which elucidated variations in the heat shock kinetics of Apg-1 and Hsp105 [66, 67]. Subsequent work supports the notion that unique interactomes exist for these homologues, reinforcing the idea of functional variation among them. For instance, Apg-1 interacts with the neurite outgrowth inhibitor Nogo-a, while Apg-2 and Hsp105 do not [68]. Furthermore, chemically-induced neuronal damage leads to distinct localizations of each Hsp110 homologue. Here, Hsp105 only localizes to intact neurons, while Apg-1 and Apg-2 localize to both damaged and intact cells [69]. Serlidaki and colleagues elucidated variations in the binding preferences of each Hsp110 homologue to different Hsp70 proteins. In their study, the authors measured the binding preference of each Hsp110 to HSPA1A and HSPA1L, two highly homologous Hsp70 proteins [63]. While each homologue preferred HSPA1A over HSPA1L, the degree of this binding preference was unique for each protein [63]. Several other pieces of evidence indicate that these homologues have significant variation from one another. While it has been observed that all three proteins are localized to the cytosol, the amount of each protein differs [69]. Moreover, our mass spectrometry results also support the notion of individual function, as no compensatory changes were observed in our Apg-1 and Apg-2 KO cell lines. Interestingly, the Apg-2 KO demonstrated a slight increase in cellular viability, while the Apg-1 and Hsp105 KOs demonstrated no alterations. These results could suggest that Apg-2 has a uniquely detrimental role in neuronal cell function. Although encouraging, these results from our study necessitate further experimental validation.

Overall, our results support the concept that Hsp110 chaperones can promote cellular prion infection through multiple complementary pathways. A wealth of literature showcases that this disease-promoting effect operates not only through Hsp70-dependent disaggregation and the generation of seeding-competent species, but also via proteostatic crosstalk – where Hsp110 expression may suppress protective autophagic clearance. Moreover, the distinct PrP^Sc^ reduction observed across individual Hsp110 knockouts highlights the salience of isoform-specific roles. Given their unique structural profiles, distinct interactomes, and differential affinities for Hsp70 proteins, individual mammalian Hsp110s likely exert specialized control over prion propagation. Collectively, these mechanisms – disaggregase function, autophagy inhibition, and autonomous chaperone activity, provide a comprehensive framework that explains why Hsp110 reduction attenuates cellular prion infection and may define novel therapeutic targets.

## Acknowledgements

### Funding

This work was supported by the National Institutes of Health (grant R01 NS121016) and Alberta Innovates (Alberta Prion Research Institute grant 201700013). We are grateful for financial support from Alberta Innovates (Alberta Prion Research Institute grant 201600023) and from t h e Office of the Chief Scientist, Alberta Environment and Protected Areas.

## Author contributions

Conceptualization, funding acquisition, project administration and supervision: HMS Supervision, project administration, formal analysis, data curation, investigation, methodology, visualization and validation: BA

Formal analysis, data curation, investigation, methodology, visualization and validation: MM, MA, CS and CMV.

Writing: HMS, BA, MM, MA.

All authors reviewed, edited, and approved the manuscript for publication. HMS is the corresponding author and assumes full responsibility for the manuscript. All authors read and approved the final version of the manuscript.

## Materials & Correspondence

All data and code needed to evaluate and reproduce the conclusions in the paper are present in the paper and/or the Supplementary Materials. Correspondence and material requests should be addressed to Dr. Hermann Schatzl.

## List of Supplementary Materials

Supplementary Information SFig1

SFig2 SFig3 SFig4

