## Supplementary data for "Distinct roles of Hsp110 proteins in regulating prion propagation"

### **Supplementary Materials**

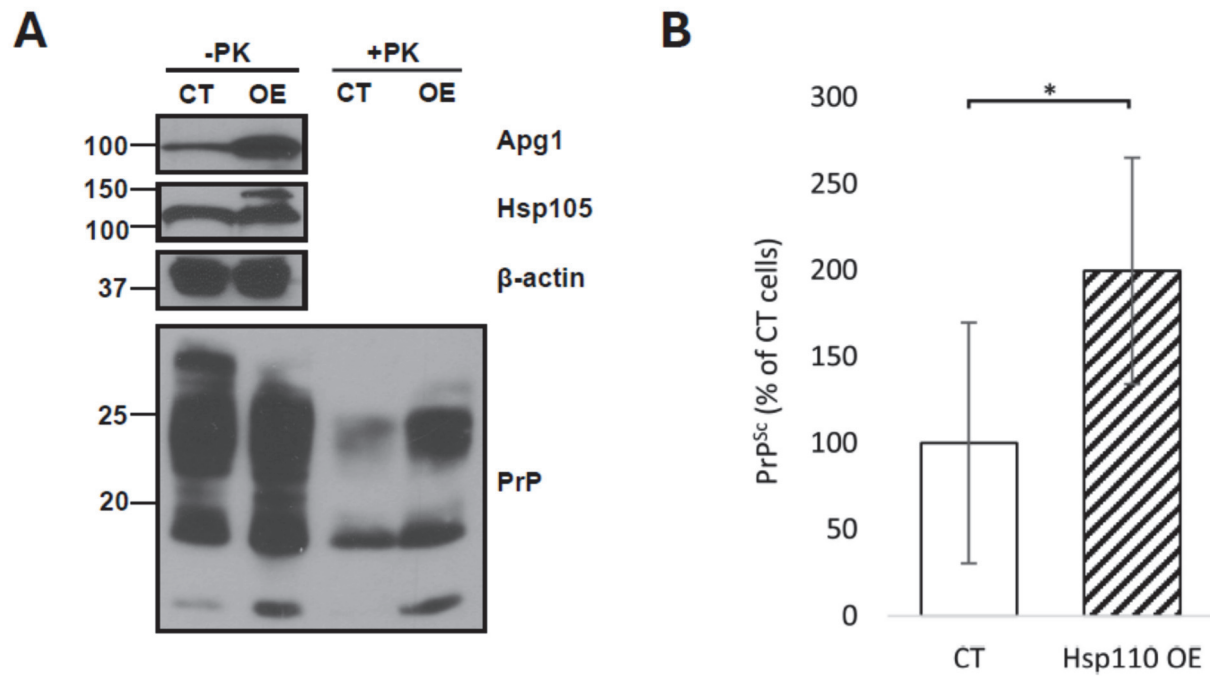

**Supplementary Fig 1. PrP<sup>Sc</sup> levels increase following transient overexpression of Apg-1 and Hsp105 in persistently prion-infected ScN2a cells.**

**(A)** Representative immunoblot illustrating increased PrP<sup>Sc</sup> levels following 96 hours of overexpression (OE) in comparison to control cells (CT). 22L-ScN2a cells were co-transfected with 2 µg of Apg-1 and Hsp105 (Hsp110 OE), or empty vector (CT), twice at 48-hour intervals. At 96 h post-initial transfection, post-nuclear lysates were collected and analyzed via immunoblotting. To elucidate PrP<sup>Sc</sup>, lysates were subjected to PK digestion (+PK) or not (-PK). PK-resistant PrP<sup>Sc</sup> (+PK) shows up as un-glycosylated (19 kDa), mono- (23/24 kDa) and di-glycosylated bands (27-30 kDa). Blots were probed with monoclonal antibodies against Apg-1, Hsp105, beta-actin and PrP. **(B)** Densitometric quantification (mean ± SD) demonstrating a significant increase in PrP<sup>Sc</sup> levels in the overexpressing cells. PrP<sup>Sc</sup> levels were quantified via densitometric analysis and normalized to the beta-actin loading control. Statistical analysis was performed using a Student's *t*-test (\* p<0.05).

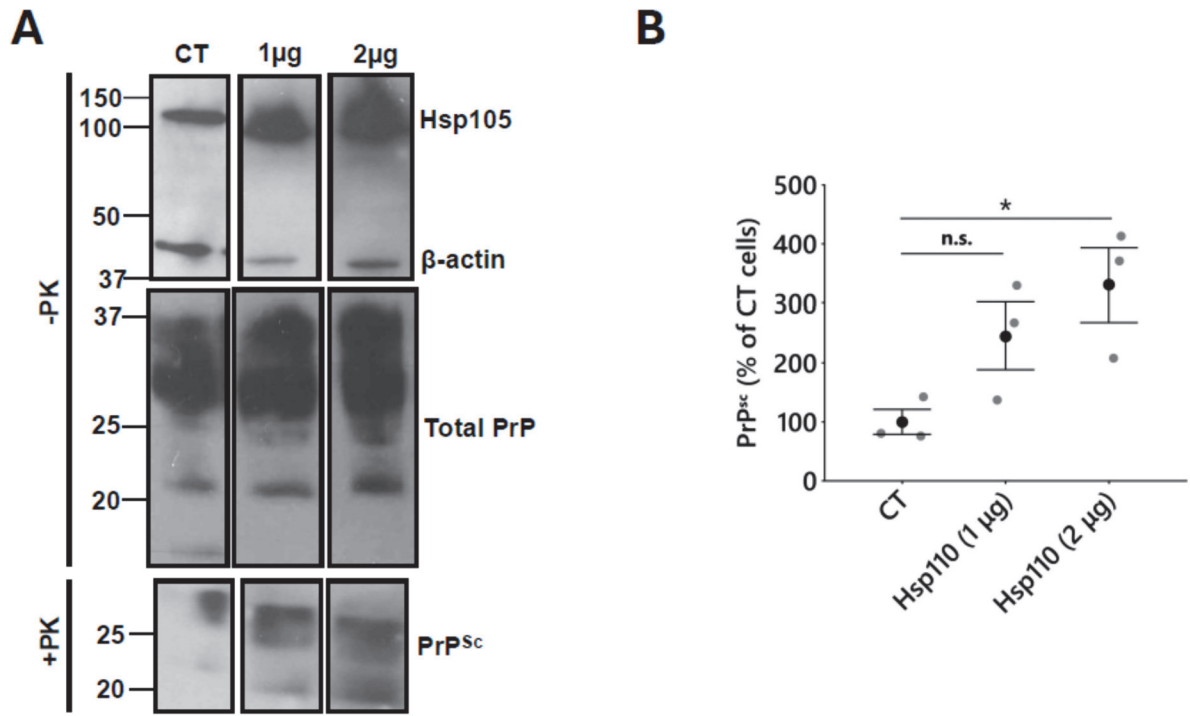

**Supplementary Fig 2. Dose-dependent increase in PrP<sup>Sc</sup> following transient overexpression of all Hsp110s in persistently prion-infected ScN2a cells.**

**(A)** Representative immunoblot demonstrating a dose-dependent increase in PrP<sup>Sc</sup> levels 96 post-overexpression. 22L-ScN2a cells were transfected with empty vector (CT), or a mixture (1 or 2 µg) of vectors encoding all three Hsp110 family members. Transfections occurred twice at 48-hour intervals, and at 96 hours post-initial transfection, post-nuclear lysates were collected and analyzed via immunoblotting. Lysates were subjected to PK digestion (+PK) or not (-PK). PK-resistant PrP<sup>Sc</sup> (+PK) shows up as unglycosylated (19 kDa), mono- (23/24 kDa) and di-glycosylated bands (27-30 kDa). Blots were probed with monoclonal antibodies against Hsp105, beta-actin and PrP (+/- PK). **(B)** Densitometric quantification (mean ± SEM) of PrP<sup>Sc</sup> levels. PrP<sup>Sc</sup> levels were quantified via densitometric analysis and normalized to the beta-actin loading control. Statistical analysis was performed using a One-way *ANOVA* with Tukey's HSD test for pair-wise comparisons ("n.s." signifies non-significance; \* *p*<0.05). A significant increase was observed in cells transfected with 2 µg plasmid DNA.

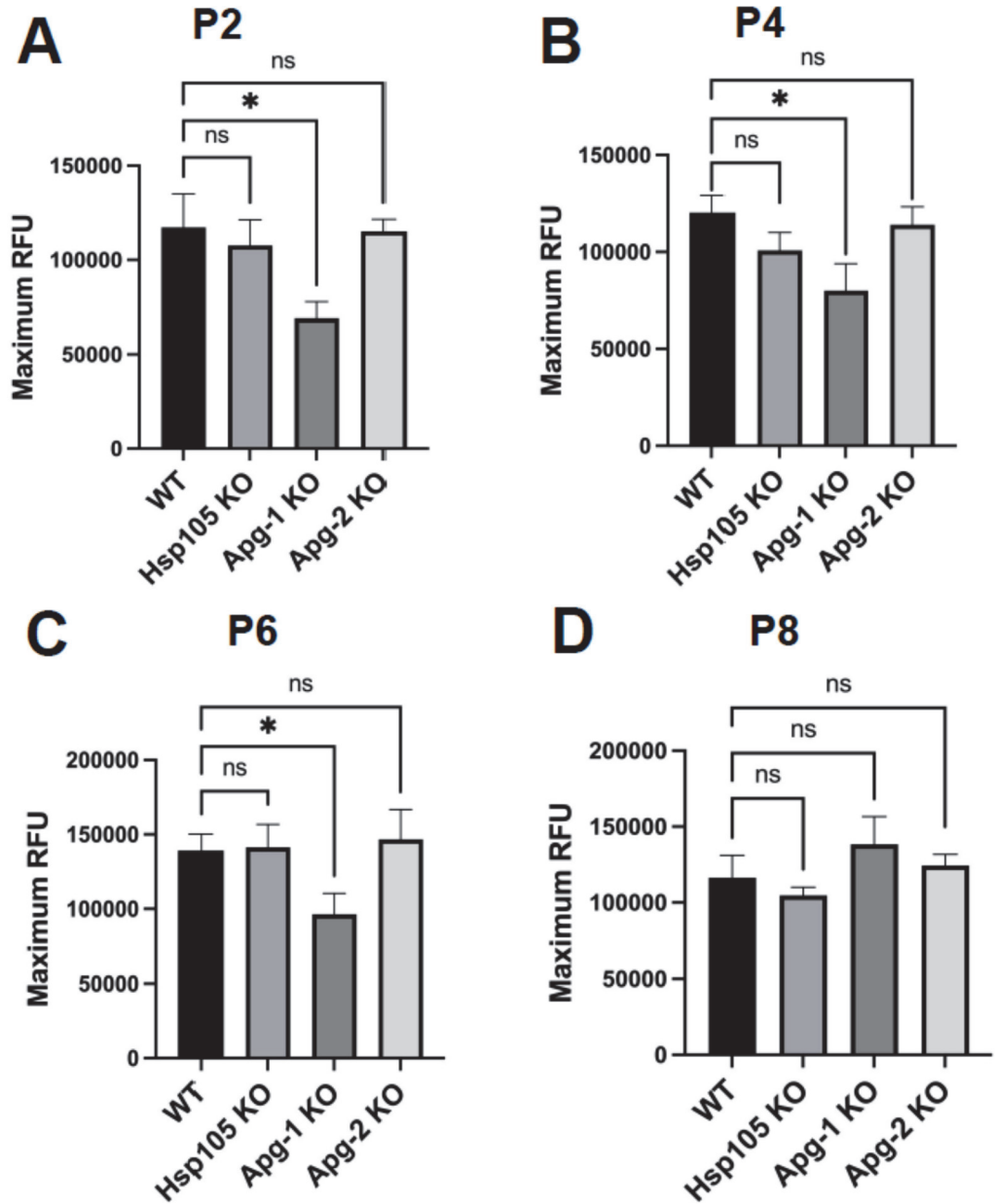

**Supplementary Fig 3. Apg-1 KO demonstrates a significant reduction in maximum fluorescence across several passages in RT-QuIC analysis.**

(A-D) Bar graphs (mean ± SEM) illustrating maximum relative fluorescence intensity (RFU) at P2 (A), P4 (B), P6 (C), and P8 (D) for each cell line. Apg-1 KO cells demonstrate significant reductions at P2, P4, and P6. The maximum fluorescence values were obtained via MARS data analysis software. Statistical analyses were performed using one-way ANOVA with a Dunnett's test for pair-wise comparisons (P2), the Kruskal-Wallis test with Dunn's test for pair-wise comparisons (P4 and P6), and Welch's ANOVA (P8) ("ns" signifies non-significance; \* $p < 0.05$). Visualized in GraphPad Prism 11.

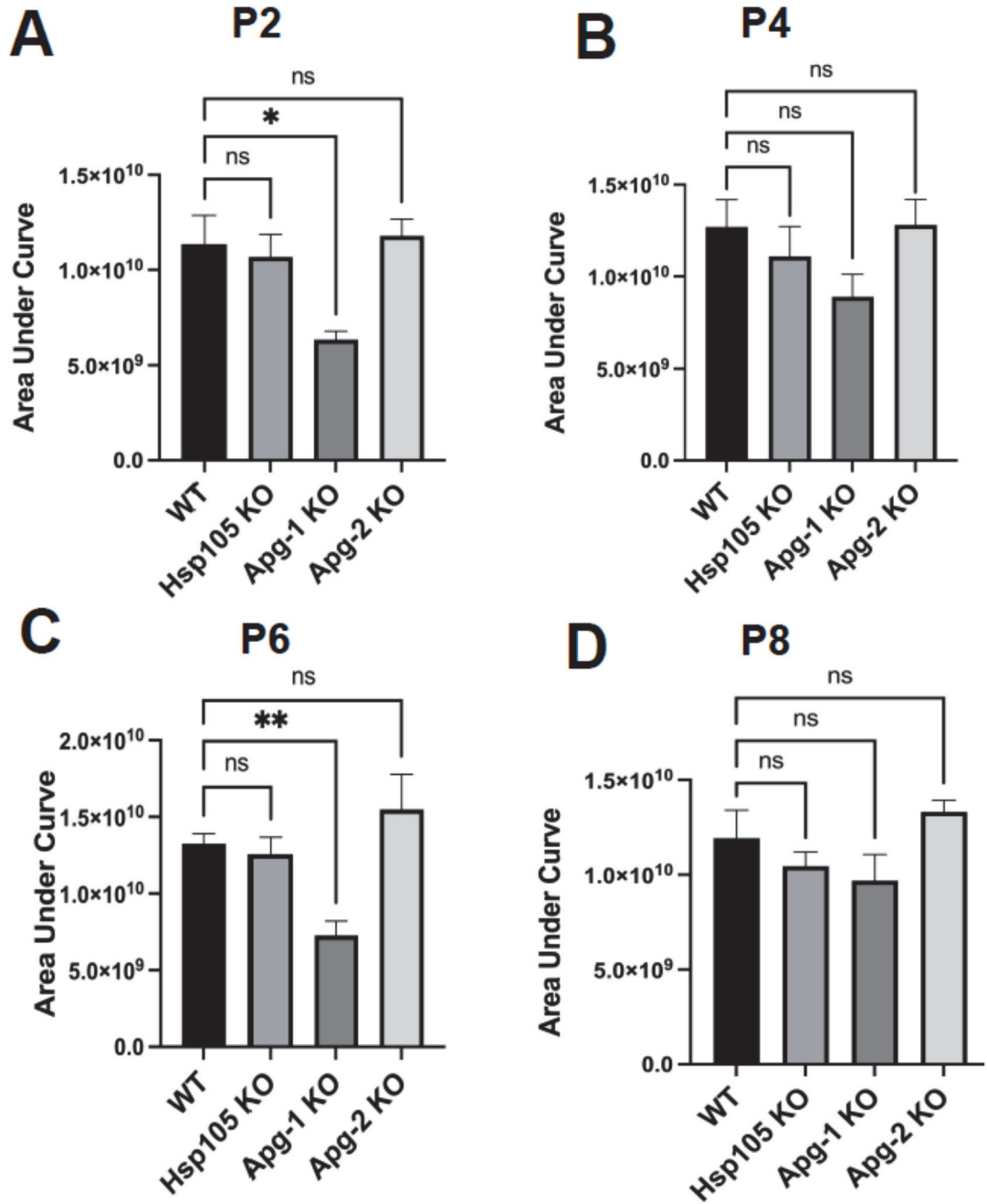

**Supplementary Fig 4. Apg-1 KO demonstrates significant area under the curve reductions at multiple passages in RT-QulC analysis.**

(A-D) Bar graphs (mean ± SEM) illustrating area under the curve for P2 (A), P4 (B), P6 (C), and P8 (D) of each cell line. Area under the curve values were obtained via MARS data analysis software. Apg-1 KO cells demonstrate significant reductions at P2 and P6, and trends at passages 4 and 8. Statistical analyses were performed using Welch's ANOVA with a Dunnett's T3 test for pair-wise comparisons (P2), the Kruskal-Wallis test with Dunn's test for pair-wise comparisons (P4 and P6), and one-way ANOVA (P8) ("ns" signifies non-significance; \* $p < 0.05$; \* $p < 0.01$). Visualized in GraphPad Prism 11.
